# The FIGNL1-interacting protein C1orf112 is synthetic lethal with PICH and mediates RAD51 retention on chromatin

**DOI:** 10.1101/2022.10.07.511242

**Authors:** Colin Stok, Nathalie van den Tempel, Marieke Everts, Elles Wierenga, Femke Bakker, Yannick Kok, Inês Teles Alves, Lucas T. Jae, Arkajyoti Bhattacharya, Elefteria Karanika, Polina Perepelkina, Steven Bergink, Kok-Lung Chan, H. Rolf de Boer, Rudolf S.N. Fehrmann, Marcel A.T.M. van Vugt

## Abstract

Joint DNA molecules are natural by-products of DNA replication and repair. Persistent joint molecules give rise to ultrafine DNA bridges (UFBs) in mitosis, which compromise sister chromatid separation. The DNA translocase PICH (*ERCC6L*) plays a central role in UFB resolution. A genome-wide loss-of-function screen was performed to identify the genetic contexts in which cells become dependent on PICH. In addition to genes involved in DNA condensation, centromere stability and DNA damage repair, we identified the uncharacterized protein C1orf112. We find that C1orf112 interacts with and stabilizes the AAA+ ATPase FIGNL1. Inactivation of either C1orf112 or FIGNL1 resulted in UFB formation, prolonged retention of RAD51 on chromatin, impaired replication fork dynamics, and consequently impaired genome maintenance. Combined, our data reveal that inactivation of C1orf112 and FIGNL1 dysregulates RAD51 dynamics at replication forks, resulting in DNA replication defects, and a dependency on PICH to preserve cell viability.

## Introduction

Safeguarding genome integrity requires faithful DNA replication, and subsequently proper sister chromatid separation. Compromised DNA maintenance results in genome instability, one of the hallmarks of cancer (Hanahan & Weinberg, 2011). In many cancers, oncogene overexpression results in erroneous orchestration of DNA replication (Kotsantis et al., 2018). Consequently, cancer cells experience perturbed replication fork progression, a state known as replication stress (Gaillard et al., 2015). Cells are equipped with dedicated pathways that ensure protection and promote restart of stalled replication forks. The replication stress response is intricately coordinated with the cell cycle to ensure that DNA replication is completed before the onset of mitosis (Jackson & Bartek, 2009). Entry into mitosis with unresolved replication intermediates may result in sister chromatid non-disjunction, and subsequent loss of genome integrity. Joint DNA molecules can arise during replication in the form of topological linkages, at sites of DNA under-replication, or when DNA repair intermediates remain unresolved before cells enter mitosis.

Persistent connections between the sister chromatids result in formation of ultrafine DNA bridges (UFBs) in anaphase. UFBs differ from conventional ‘bulky’ chromatin bridges in their absence of histones, and their inability to be stained with DNA dyes, such as DAPI (Chan et al., 2007). UFBs can be induced by various factors, including topological linkages, such as DNA catenanes, that are particularly prevalent at centromeres. In cells suffering from replication stress, DNA underreplication is a major source of persistent joint DNA molecules. Especially difficult-to-replicate regions, such as common fragile sites (CFSs), remain largely under-replicated until late-S or G_2_-phase (Li & Wu, 2020). Cells that enter mitosis with incompletely replicated DNA, initiate POLD3-dependent mitotic DNA synthesis (MiDAS) (Bhowmick et al., 2016; Minocherhomji et al., 2015). MiDAS marks a final opportunity for cells to finish DNA synthesis during prophase and prometaphase. Failure to correctly complete DNA replication, results in formation of UFBs that are flanked by FANCD2 foci, which are not present at centromeric UFBs (K. L. Chan et al., 2009). UFBs flanked by FANCD2 frequently form at CFSs and other difficult-to-replicate regions (K. L. Chan et al., 2009).

Joint DNA molecules may also arise when DNA repair intermediates, such as Holliday junctions, are not resolved before mitotic entry. To prevent these late-stage repair intermediates from interfering with chromosome segregation, several nucleases are hyperactivated in the G2/M phase of the cell cycle. Specifically, mitotic kinases CDK1 and PLK1 stimulate activity of SLX1, MUS81-EME1 and XPF-ERCC1 structure-selective endonucleases, which cleave joint DNA molecules in the G2/M-phase of the cell cycle (Wyatt et al., 2013, 2017). Additionally, the cytoplasmic nuclease GEN1 acquires access to joint DNA molecules upon mitotic nuclear membrane disassembly (Y. W. Chan & West, 2014). Hyperactivation of DNA repair by homologous recombination (HR), due to loss of 53BP1 or combined loss of GEN1 and MUS81, results in an increase in joint DNA molecules and consequently formation of UFBs (Y. W. Chan et al., 2018; Tiwari et al., 2018).

The SNF2 family DNA translocase PICH, encoded by the *ERCC6L* gene, is rapidly recruited to UFBs (C. Baumann et al., 2007). PICH has high affinity for DNA under tension, and promotes the recruitment of BLM, TOP3A and RIF1 to UFBs (Broderick et al., 2015; Chan et al., 2007; Germann et al., 2014; Hengeveld et al., 2015; Ke et al., 2011). Disruption of the *Ercc6l* gene in mice leads to embryonic lethality (Albers et al., 2018), implying that PICH is required during early development. Gene expression analysis revealed that *ERCC6L* expression is increased in a range of tumour types (Pu et al., 2017). However, it is currently unclear to what extent tumour cells are dependent on PICH-mediated UFB resolution to maintain genome stability and cell viability. Here we present a genetic loss-of-function screen to identify synthetic lethal interactions with PICH, to uncover the genetic contexts in which cells become dependent on PICH.

## Results

### ERCC6L is essential for cell viability in a subset of tumour cell lines

The *ERCC6L* locus was mutated in human haploid HAP1 cells using CRISPR/Cas9-mediated gene editing **(Fig. 1A)**. Inactivation of *ERCC6L (ERCC6L^KO^*) did not significantly affect doubling time, colony forming capacity or cell cycle distribution of HAP1 cells **(Fig. 1B, C, Suppl. Fig. 1A, B)**. Similarly, introduction of short hairpin RNAs (shRNAs) targeting *ERCC6L* did not significantly inhibit cell growth of MCF-7 and MDA-MB-231 breast cancer cell lines (**Suppl. Fig. 1C-E)**. Moreover, no significant differences in chromosome numbers and copy number alterations (CNAs) were observed when comparing HAP1-*ERCC6L^KO^* with HAP1-*ERCC6L^WT^* cells **(Fig. 1D, Suppl. Fig. 1F)**. Overall, these results demonstrate that in this set of cancer cell lines, *ERCC6L* expression is not required for maintaining cell viability and genome stability.

**Figure 1:**
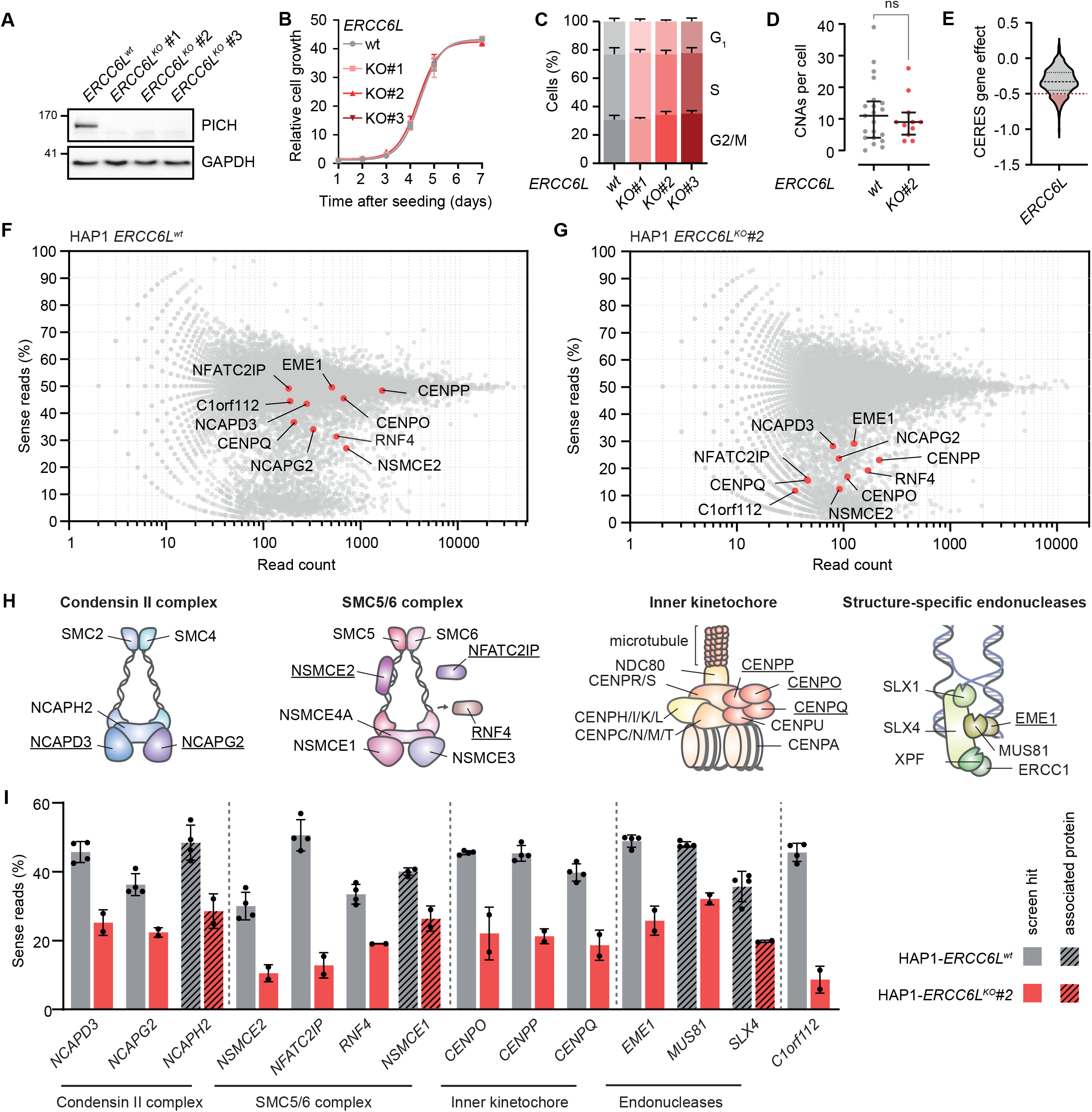
A haploid genetic screen identifies synthetic lethal genetic interactions with PICH (*ERCC6L*). **(A)** PICH protein expression levels were analysed in whole-cell lysates from three HAP1-*ERCC6L^KO^* clones by western blot. **(B)** HAP1-*ERCC6L^WT^* and HAP1-*ERCC6L^KO^* cells were seeded in 12-well plates, and cell growth was monitored for 7 days using SRB staining. Data points represent the mean and standard error of the mean (SEM) of four replicates. Curves are non-linear fits with R^2^ > 0.97. **(C)** Cell cycle profiles of HAP1-*ERCC6L^WT^* and HAP1-*ERCC6L^KO^* cells were generated by flow cytometry analysis of EdU and PI signal. Bars represent mean and standard error of the mean (SEM) of three replicates. **(D)** Copy number alterations (CNAs) were analysed using single-cell sequencing of 21 HAP1-*ERCC6L^WT^* and 11 HAP1-*ERCC6L^KO^* libraries. Statistics were performed using a two-tailed Mann-Whitney test. **(E)** *ERCC6L* CERES-corrected gene essentiality scores for 582 cell lines were obtained from the Depmap consortium. Cell lines with CERES scores below −0.5 were considered PICH-dependent. **(F-G)** Sense and anti-sense gene-trap insertions were mapped to the human genome, and a reduction in the percentage of sense insertions was used as a measure of gene essentiality. One representative replicate of a HAP1-*ERCC6L^WT^* dataset and one representative replicate of a HAP1-*ERCC6L^KO^* are presented. Synthetic lethal genes were selected based on two-sided Fisher’s exact tests. **(H)** Identified synthetic lethal genes with *ERCC6L* were categorised in four protein complexes: the condensin II complex (*NCAPD3, NCAPG2*), the SMC5/6 complex (*NSMCE2, NFATC2IP, RNF4*), the inner kinetochore complex (*CENPO, CENPP, CENPQ*), and the SLX4 tri-nuclease complex (*EME1*). **(I)** Percentage of sense insertions for the genes identified as synthetic lethal *ERCC6L* interactors, and for complex members. Datapoints represent data from four independent HAP1-*ERCC6L^WT^* datasets and two independent HAP1-*ERCC6L^KO^* datasets.

To extend our analysis to other cell line models, we analysed *ERCC6L* perturbation effects in the Cancer Dependency Map (DepMap) (Dempster et al., 2019; Tsherniak et al., 2017). *ERCC6L* was scored as ‘essential’ in 96 out of 582 cancer cell lines (from here on referred to as “*ERCC6L*-dependent” cell lines) **(Fig. 1E, Suppl. Fig. 1G)**. Conversely, the majority of the cancer cell lines can be classified as “*ERCC6L*-independent”. No apparent correlations were found between tissue type of origin and *ERCC6L* gene effect score **(Suppl. Fig. 1G)**, although *ERCC6L* was not essential in any of the leukaemia cell lines. Using expression data from the Cancer Cell Line Encyclopedia (CCLE), no differences were observed between *ERCC6L* mRNA expression levels in *ERCC6L*-dependent and *ERCC6L*-independent cell lines **(Suppl. Fig. 1H)**. Moreover, the number of genome-wide copy number alterations, used as a proxy for genomic instability, was similar in *ERCC6L*-dependent and *ERCC6L*-independent cell lines **(Suppl. Fig. 1I)**. To better understand the genetic context determining dependency on *ERCC6L*, gene set enrichment analysis (GSEA) was performed on mRNA expression data of *ERCC6L*-dependent and *ERCC6L*-independent cell lines **(Suppl. Fig. 1J, Suppl. Data 1)**. Genes involved in DNA repair and DNA recombination were generally expressed at lower levels in *ERCC6L*-dependent cells **(Suppl. Fig. 1J)**, suggesting that *ERCC6L* becomes more important for cell viability in cells with relatively low expression of DNA maintenance genes. Interestingly, gene sets related to the adaptive immune response were also depleted in *ERCC6L*-dependent cells, possibly due to the absence of leukaemia cell lines in the *ERCC6L*-dependent panel of cell lines **(Suppl. Fig. 1G)**.

### A loss□of□function screen identifies synthetic lethal interactions with ERCC6L

To gain further insight into the genetic context of *ERCC6L*-dependency, a genome-wide loss□of□function insertional mutagenesis screen was performed to identify essential genes in HAP1-*ERCC6L^KO^* cells **(Suppl. Data 2)**. Common essential genes were filtered out using reference insertional mutagenesis screens in parental HAP1 cells **(Fig. 1F)** (Blomen et al., 2015). Combined analysis of two independent screens identified ten genes that specifically affected the viability of HAP1-*ERCC6L^KO^* cells **(Fig. 1G)**. The identified screen hits included genes involved in resolution of joint DNA molecules (*EME1*) (Abraham et al., 2003), kinetochore organization (*CENPO, CENPP, CENPQ*) (Hori et al., 2008), DNA condensation (*NCAPD3, NCAPG2*) (Ono et al., 2003), and processing of collapsed replication forks (*NSMCE2*) (Pond et al., 2019) **(Fig. 1G-H)**. Moreover, *NFATC2IP*, the human ortholog of *Schizosaccharomyces pombe Rad60* (Morishita et al., 2002), was identified as one of the hits **(Fig. 1G)**, which has been described to promote the function of NSMCE2 in DNA repair (Prudden et al., 2011). Mutations in some of these genes, including *NCAPD3, NSMCE2* and *EME1*, have previously been associated with increased numbers of UFBs (Martin et al., 2016; Pond et al., 2019; Ying et al., 2013), showing that the screening set-up identified genes that are relevant in the context of UFB biology. Genes from the same protein complexes as the screen hits generally showed a trend toward synthetic lethality with *ERCC6L* as well, including *MUS81, NCAPH2* and *NSMCE1* **(Fig. 1H-I)**. Additionally, we identified a synthetic lethal interaction between the uncharacterized open reading frame *C1orf112* and *ERCC6L* **(Fig. 1G)**. As a general screen validation, shRNAs targeting *NCAPD3, NFATC2IP* and *C1orf112* were introduced in HAP1-*ERCC6L^WT^* cells and HAP1-*ERCC6L^KO^* cells **(Suppl. Fig. 1K)**. Subsequently, HAP1-*ERCC6L^WT^* cells and HAP1-*ERCC6L^KO^* cells were mixed in a 1:1 ratio, and the *ERCC6L* mutation was monitored over time using TIDE (Brinkman et al., 2014). Upon depletion of *NCAPD3, NFATC2IP* and *C1orf112*, the number of cells with the *ERCC6L* mutation decreased rapidly over time **(Suppl. Fig. 1L)**, confirming the synthetic lethality between these genes. Due to its enigmatic nature, we focussed on exploring the function of the *C1orf112* gene, and its relation to *ERCC6L*.

### C1orf112 is predicted to function in DNA maintenance

Function predictions were made for C1orf112 using our previously established ‘GenetICA’ guilt-by-association platform (Urzúa-Traslaviña et al., 2021) **(Fig. 2A, Suppl. Data 3)**. Based on similarity in mRNA expression patterns, *C1orf112* was predicted to be involved in DNA repair, DNA replication, and sister chromatid segregation **(Fig. 2A)**. *C1orf112* shows particularly similar mRNA expression patterns to key HR genes, such as BRCA1, BRCA2 and RAD51 **(Fig. 2A)**. Indeed, Gene network analysis shows tight clustering of C1orf112 within the cloud of GO_DNA_REPAIR genes **(Fig. 2B)**. Overall, these data suggest that the C1orf112 protein may play a role in DNA repair or recombination.

**Figure 2:**
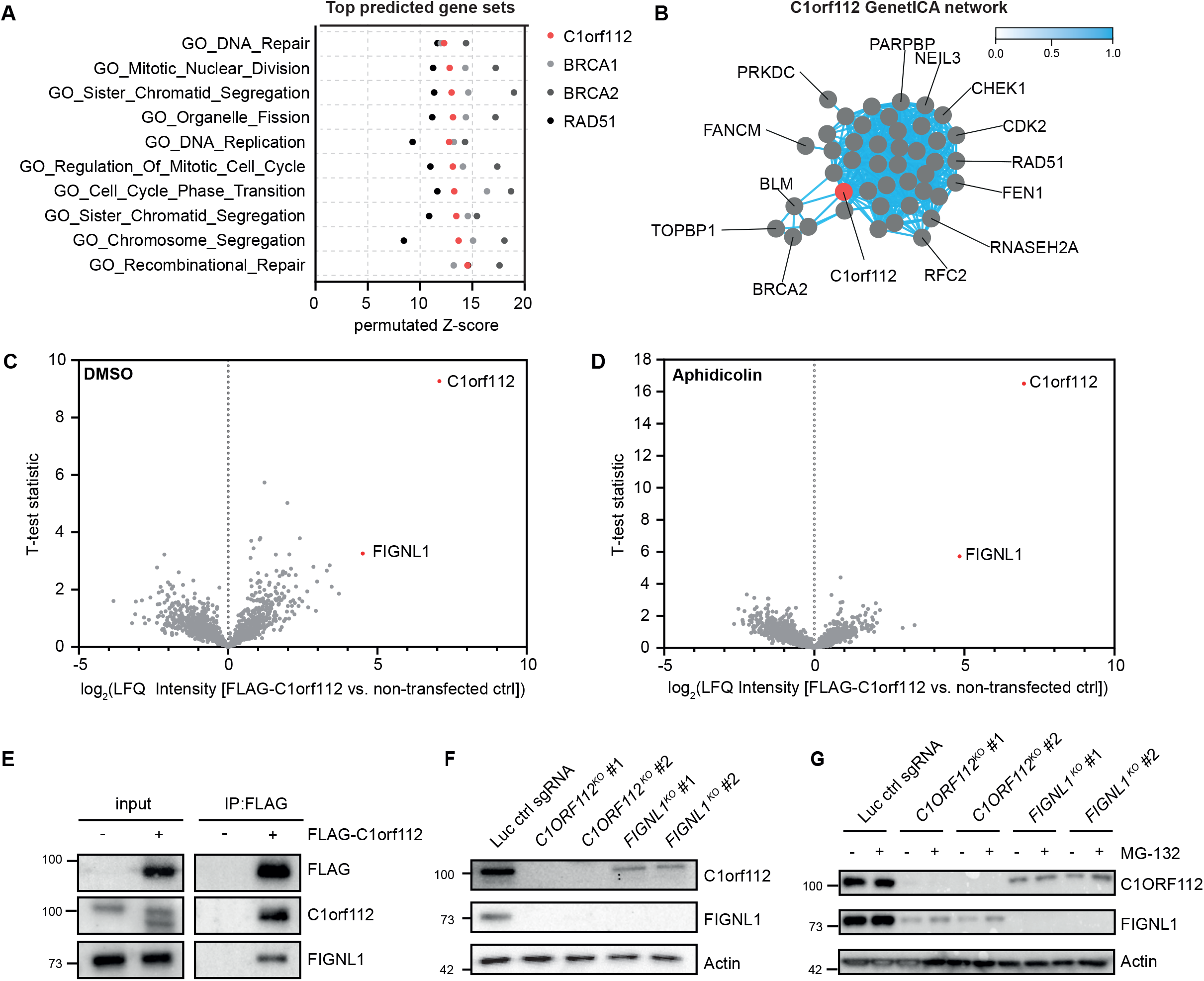
C1orf112 interacts with the AAA+ ATPase FIGNL1. **(A)** Function predictions for C1orf112 were performed using permutated Z-scores using GenetICA. Ten highest ranking gene sets from ‘Gene Ontology - Biological Process’ are plotted. Permutated Z-scores were compared to those for key HR proteins BRCA1, BRCA2 and RAD51. **(B)** A co-functionality network was plotted using C1orf112 and the top genes from the GO_DNA_REPAIR gene set. Blue lines indicate strong coexpression profiles. **(C)** HEK293T cells were transiently transfected with FLAG-C1orf112, and cells were treated with DMSO or 0.2 μM aphidicolin (APH) for 24 h. After anti-FLAG M2 immunoprecipitation (IP), proteins were digested in-gel, and analysed by HPLC-MS. Graphs represent mean log_2_(LFQ Intensity) scores of three biological replicates, p-values were calculated by t-tests. **(E)** HEK293T cells were transfected with FLAG-C1orf112, and IP reactions were performed in whole-cell lysates using anti-FLAG M2 beads. C1orf112 and FIGNL1 levels in input and IP fraction were analysed by western blot. **(F)** C1orf112 and FIGNL1 protein expression levels were analysed in whole-cell lysates from two HAP1-*C1orf112^KO^* clones and two HAP1-*FIGNL1^KO^* clones by western blot. **(G)** HAP1-*C1orf112^KO^* and HAP1-*FIGNL1^KO^* cells were treated with 10 μM MG-132 proteasome inhibitor for 3 h. C1orf112 and FIGNL1 protein expression levels were analysed in whole-cell lysates by western blot.

### C1orf112 interacts with the AAA+ ATPase FIGNL1

To further investigate a possible function of C1orf112, we aimed to express FLAG-tagged C1orf112 in HEK293T cells. A transcript variant analysis was performed to identify which C1orf112 isoform was predominantly expressed. The *C1orf112* locus encompasses a region of 192,073 nucleotides at chromosome position 1q24.2, encoding 32 putative exons that can be spliced into five protein-coding transcripts and four non-coding transcripts **(Suppl. Fig. 2A)**. Pan-tissue expression of the different transcript variants was analysed using RNA-seq data from healthy tissue samples from the Genotype-Tissue Expression (GTEx) project, revealing the expression of two protein-coding transcripts (ENST00000359326, ENST00000496973) and one non-coding transcript (ENST00000459772) **(Suppl. Fig. 2B)**. The expression of these three transcripts was validated in HeLa and RPE-1 cells by RT-qPCR using primers spanning exon-exon boundaries unique to each transcript **(Suppl. Fig. 2C)**. On the protein level, no evidence was found for expression of the shorter ENST00000496973 isoform (data not shown), and we therefore focussed on the longer protein-coding *C1orf112* transcript ENST00000359326.

FLAG-tagged C1orf112 was transiently expressed in HEK293T cells, followed by immunoprecipitation (IP) and mass spectrometry (MS) to identify C1orf112-interacting proteins **(Fig. 2C, D, Suppl. Fig. 2D, Suppl. Data 4)**. Strong enrichment was observed for peptides mapping to the AAA+ ATPase FIGNL1, identifying FIGNL1 as a potential interactor of C1orf112 **(Fig. 2C, D)**. The interaction between FLAG-tagged C1orf112 and endogenous FIGNL1 was validated by immunoblotting **(Fig. 2E)**. The interaction between C1orf112 and FIGNL1 was also observed in conditions of aphidicolin-induced replication stress **(Fig. 2E)**, upon induction of DNA damage by cisplatin treatment **(Suppl. Fig. 2D)**, and upon enrichment of mitotic cells using nocodazole **(Suppl. Fig. 2D)**. Conversely, endogenous C1orf112 was identified as an interactor of FLAG-tagged FIGNL1 by mass spectrometry, and validated by immunoblotting **(Suppl. Fig. 2D, E)**. Although *FIGNL1* did not reach significance based on the threshold in our loss-of-function screen, closer inspection of the data did reveal a reduction of FIGNL1-inactiving mutations in HAP1 *ERCC6L^KO^* cells **(Suppl. Fig. 2F)**.

To further explore the function of C1orf112 and FIGNL1, HAP1-*C1orf112^KO^* and HAP1-*FIGNL1^KO^* cells lines were generated using CRISPR/Cas9 **(Fig. 2F)**. Inactivation of C1orf112 strongly reduced protein levels of FIGNL1 **(Fig. 2F)**. FIGNL1 inactivation also led to reduced C1orf112 levels, albeit less pronounced **(Fig. 2F)**. Proteasome inhibition using MG-132 partially restored FIGNL1 expression levels in *C1orf112^KO^* cells **(Fig. 2G, Suppl. Fig. 2G)**, suggesting that C1orf11 stabilizes FIGNL1 by protecting it from proteasomal degradation. Of note, MG-132 treatment did not rescue C1orf112 protein levels in *FIGNL1^KO^* cells, which may indicate an alternative turnover mechanism **(Fig. 2G, Suppl. Fig. 2G)**. Interestingly, GenetICA protein function predictions for FIGNL1 were highly similar to those for C1orf112, with both proteins being predicted to function in DNA repair **(Suppl. Fig. 2H)**. Moreover, cell lines in the DepMap dataset that strongly depend on C1orf112 for their survival, also frequently relied on FIGNL1, indicating strong co-dependence **(Suppl. Fig. 2I)**. Taken together, the observed protein-protein interaction, co-depletion, predicted co-functionality, and co-dependency observed for C1orf112 and FIGNL1 suggest that the two proteins function together in one protein complex.

### Inactivation of C1orf112 or FIGNL1 sensitizes cells to DNA damaging agents

To investigate a potential role for C1orf112 in DNA damage repair, HAP1-*C1orf112^KO^* and HAP1-*FIGNL1^KO^* cells were treated with a panel of DNA damaging agents, and cell viability was monitored using MTT assays and colony formation assays, using HAP1-*FANCD2^KO^* cells as a positive control **(Fig. 3A, Suppl. Fig 3A, B)**. HAP1-*C1orf112^KO^* and HAP1-*FIGNL1^KO^* cells displayed increased sensitivity to crosslinking agents, including cisplatin and mitomycin C, albeit more moderate than HAP1-*FANCD2^KO^* cells **(Fig. 3A, Suppl. Fig 3B)**. Moreover, HAP1 *C1orf112^KO^* and *FIGNL1^KO^* showed enhanced sensitivity to the replication perturbing agent hydroxyurea **(Suppl. Fig 3B)**. In contrast, no clear increase in sensitivity was observed towards the PARP inhibitor olaparib **(Fig. 3A, Suppl. Fig 3B)**. Sensitization to the DNA alkylating agent 1-methyl-3-nitro-1-nitrosoguanidine (MNNG) **(Suppl. Fig 3A)**, and the DSB-inducing agent bleomycin **(Suppl. Fig 3B)**, suggested that C1orf112 and FIGNL1 may not be involved in a single DNA repair pathway. The different degrees of sensitization to DNA damaging agents between HAP1-*C1orf112^KO^* cells and HAP1-*FANCD2^KO^* cells, suggested that C1orf112 is not part of the canonical DNA cross-link repair pathway. Moreover, FANCD2 mono-ubiquitination upon cisplatin treatment, a read-out of Fanconi anaemia pathway functionality (Walden & Deans, 2014), was not affected upon *C1orf112* or *FIGNL1* inactivation **(Suppl. Fig 3C)**.

**Figure 3:**
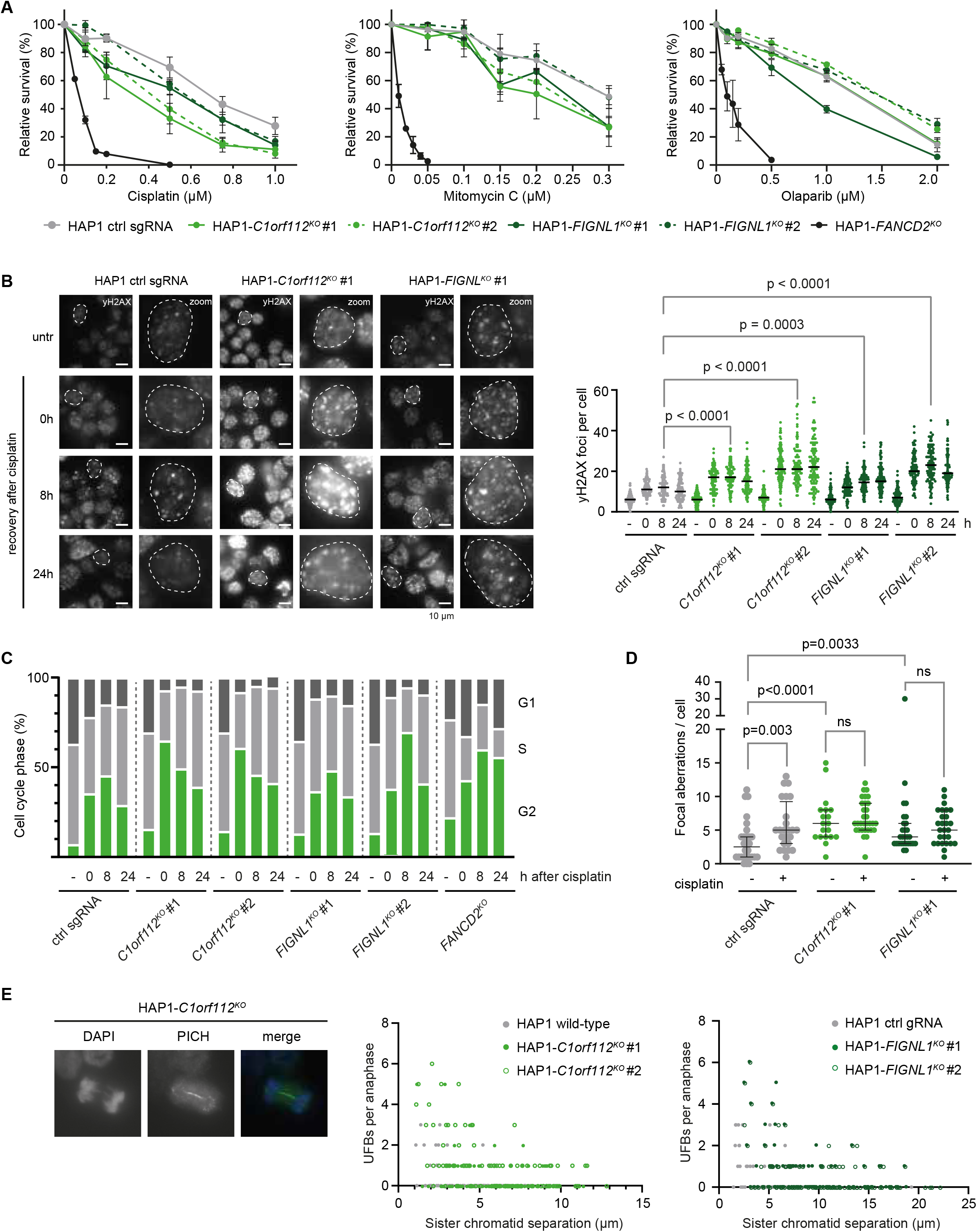
Inactivation of C1orf112 or FIGNL1 is associated with sensitivity to DNA damaging agents and accumulation of DNA damage. **(A)** HAP1-sgLUC, HAP1-*C1orf112^KO^*, HAP1-*FIGNL1^KO^* and HAP1-*FANCD2^KO^* cells were treated with cisplatin for 24 h, mitomycin C for 24 h or olaparib for 7 days. Colony formation was measured after 7 days by Coomassie Brilliant Blue staining. Graphs represent mean and standard error of the mean (SEM) of three independent biological replicates. **(B)** HAP1-sgLUC, HAP1-*C1orf112^KO^*, HAP1-*FIGNL1^KO^* and HAP1-*FANCD2^KO^* cells were treated with 1 μM cisplatin for 24 h, and cells were fixed at 0 h, 8 h and 24 h after washout of cisplatin. Cells were stained for γH2AX foci formation, and at least 100 cells per condition were scored. Statistics were performed using two-tailed Mann-Whitney tests. **(C)** Cell cycle profiles of HAP1-sgLUC, HAP1-*C1orf112^KO^*, HAP1-*FIGNL1^KO^* and HAP1-*FANCD2^KO^* cells were generated by flow cytometry analysis of EdU and PI signal. Bars represent means of three replicates. **(D)** Copy number alterations (CNAs) were analysed using single-cell sequencing of 28 HAP1, 19 HAP1-*C1orf112^KO^* and 28 HAP1-*FIGNL1^KO^* libraries. Cisplatin treatment was performed with 1 μM cisplatin for 96 h. CNA analysis in cisplatin-treated cells was performed using 24 HAP1, 31 HAP1-*C1orf112^KO^* and 27 HAP1-*FIGNL1^KO^* libraries. Statistics were performed using two-tailed Mann-Whitney tests. **(E)** PICH-positive anaphase bridges were analysed in unchallenged HAP1-sgLUC, HAP1-*C1orf112^KO^* and HAP1-*FIGNL1^KO^* cells. At least 60 anaphases were scored per condition.

To test whether HAP1-*C1orf112^KO^* and HAP1-*FIGNL1^KO^* cells displayed impaired DNA repair, γH2AX foci and KAP1 phosphorylation were monitored over time after cisplatin treatment **(Fig 3B, Suppl. Fig 3D)**. Directly after cisplatin treatment, and after drug wash-out, DNA damage levels were exacerbated in HAP1-*C1orf112^KO^* and HAP1-*FIGNL1^KO^* cells compared to control cells **(Fig 3B, Suppl. Fig 3D)**. Furthermore, HAP1-*C1orf112^KO^* and HAP1-*FIGNL1^KO^* cells accumulated in G_2_-phase **(Fig. 3C)**, possibly due to accumulation of unresolved DNA lesions. Additionally, HAP1-*C1orf112^KO^* and HAP1-*FIGNL1^KO^* cells displayed increased copy number alterations, as determined by single cell sequencing, already in unperturbed conditions **(Fig. 3D, Suppl. Fig 3E)**. Finally, an increase in the amount of persistent UFBs was observed in HAP1-*C1orf112^KO^* and HAP1-*FIGNL1^KO^* cells **(Fig. 3E)**. Together, these data suggest that inactivation of C1orf112 or FIGNL1 results in increased genome instability, impaired processing of cisplatin-induced DNA lesions, and G2 cell cycle arrest.

### Inactivation of C1orf112 or FIGNL1 leads to persistent chromatin association of RAD51

Because C1orf112 and FIGNL1 inactivation did not perturb the core Fanconi anaemia pathway, we next tested their involvement in the HR repair by monitoring RAD51 foci **(Fig. 4A)**. HAP1-*C1orf112^KO^* and HAP1-*FIGNL1^KO^* cells showed a strongly increased number of RAD51 foci, both in untreated conditions and after cisplatin treatment **(Fig. 4A)**. Additionally, both RAD51 foci intensity and overall nuclear fluorescence intensity were greater in HAP1-*C1orf112^KO^* and HAP1-*FIGNL1^KO^* cells **(Fig. 4A, Suppl. Fig 4A)**. In line with these observations, RAD51 was increasingly associated in the nuclear soluble and chromatin fraction of HAP1-*C1orf112^KO^* cells **(Fig. 4B)**. Fluorescence recovery after photobleaching (FRAP) assays confirmed that GFP-RAD51 dynamics were altered in HAP1-*C1orf112^KO^* cells, showing increased mobility of total nuclear GFP-RAD51 in unchallenged and cisplatin-treated conditions, despite the increased association of RAD51 with chromatin **(Suppl. Fig 4B)**, consistent with previous reports on RAD51 dynamics during replication stress (Yu et al., 2003).

**Figure 4:**
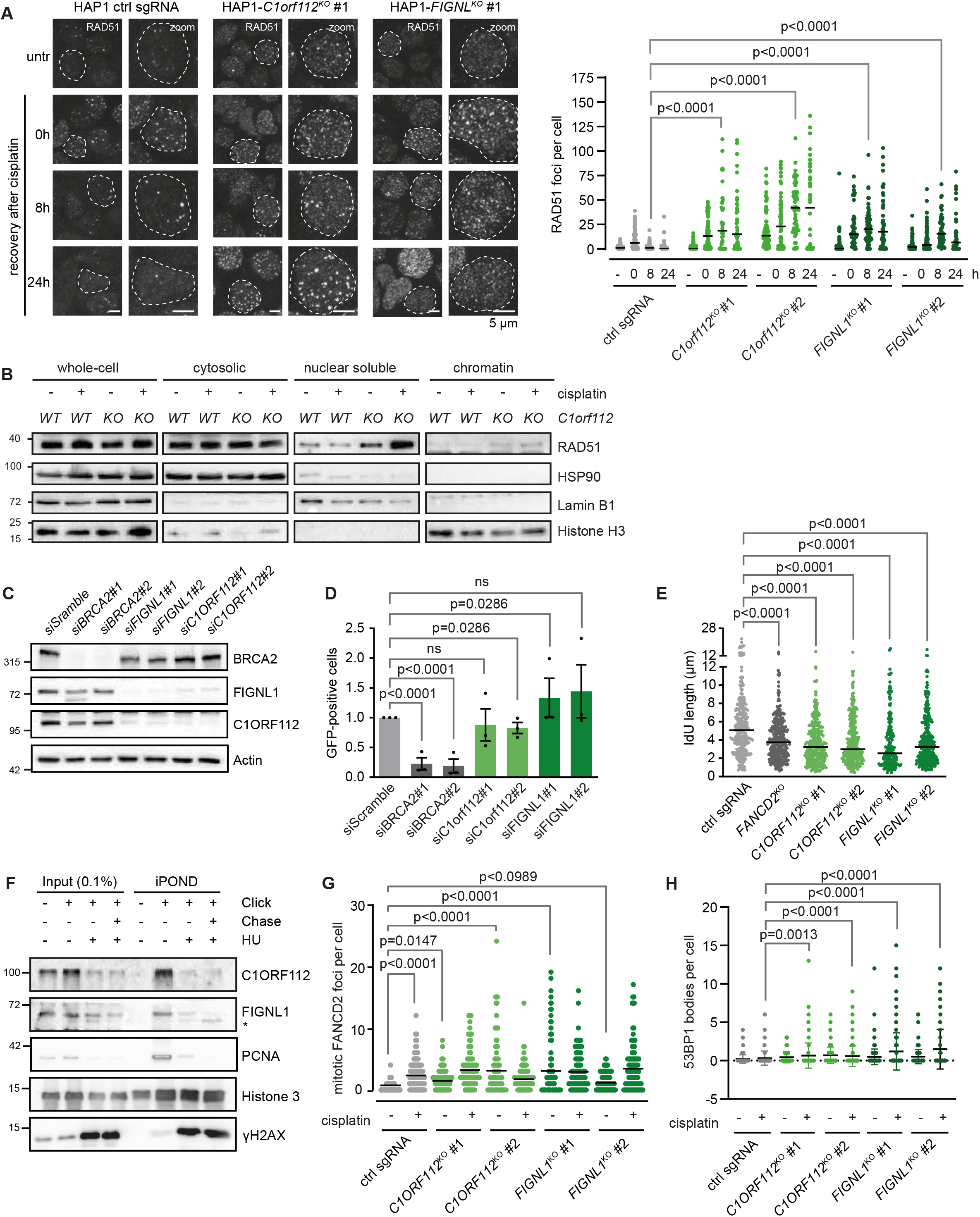
C1orf112 and FIGNL1-deficient cells display altered RAD51 and replication fork dynamics. **(A)** HAP1-sgLUC, HAP1-*C1orf112^KO^*, HAP1-*FIGNL1^KO^* and HAP1-*FANCD2^KO^* cells were treated with 1 μM cisplatin for 24 h, and cells were at fixed 0 h, 8 h and 24 h after washout of cisplatin. Cells were stained for RAD51, and at least 100 cells per condition were scored for foci formation. Statistics were performed using two-tailed Mann-Whitney tests. **(B)** HAP1-sgLUC, HAP1-*C1orf112^KO^* and HAP1-*FIGNL1^KO^* cells were treated with 1 μM cisplatin for 24 h, and whole-cell lysates, cytosolic, nuclear and chromatin protein fractions were subsequently isolated. Expression of RAD51, HSP90 (cytosolic marker), LaminB1 (nuclear marker), and Histone H3 (chromatin marker) were analysed by western blot. **(C)** HeLa DR-GFP cells were transfected with siRNAs targeting BRCA2, C1orf112 or FIGNL1. 48 h after transection, expression of BRCA2, C1orf112 and FIGNL1 was analysed by western blot. **(D)** HeLa DR-GFP cells were sequentially transfected with the indicated siRNAs and I-SceI plasmid. 48 h after I-SceI transfection, GFP-positivity was analysed by flow cytometry. Data points represent three independent biological replicates and standard error of the mean (SEM). Statistics were performed by Mann-Whitney tests. **(E)** HAP1-sgLUC, HAP1-*C1ORF112^KO^*, HAP1-*FIGNL1^KO^* and HAP1-*FANCD2^KO^* cells were pulse-labelled with IdU, followed by and CldU in the presence of 4 mM HU. Lengths of at least 200 DNA fibres were measured per condition. Statistics were performed by two-tailed Mann-Whitney tests. **(F)** HAP1 cells were labeled with EdU for 10 minutes. For chase conditions, EdU labeling was followed with thymidine treatment for 1 h. Fork stalling was induced by treatment with 4 mM HU for 2 h. Proteins at nascent DNA were cross-linked using formaldehyde, EdU was coupled to biotin-conjugated azide, and protein capture was performed using streptavidin beads. C1orf112, FIGNL1, PCNA, Histone H3 and γH2AX levels in input and iPOND fractions were analysed by western blot. **(G)** HAP1-sgLUC, HAP1-*C1ORF112^KO^* and HAP1-*FIGNL1^KO^* cells were treated with 1 μM cisplatin for 24 h, and subsequently FANCD2 foci were scored in at least 30 mitotic cells were analysed in three replicates. Statistics were performed by one-way Anova. **(H)** HAP1-sgLUC, HAP1-*C1ORF112^KO^* and HAP1-*FIGNL1^KO^* cells were treated with 1 μM cisplatin for 24 h, and subsequently 53BP1 bodies were analysed in at least 250 cells per condition per replicate. Statistics were performed using one-way Anova with Sidak’s post hoc test.

To test the consequences of altered RAD51 dynamics on HR efficacy, DR-GFP assays were performed in HeLa cells treated with siRNAs targeting *C1orf112, FIGNL1* or *BRCA2* **(Fig. 4C, D)**. Surprisingly, I-Sce1-induced gene conversion was not negatively affected by depletion of C1orf112 or FIGNL1 **(Fig. 4D)**, suggesting that HR efficiency is not diminished in these cells. Accordingly, sister chromatid exchange (SCE) assays did not show a reduction in recombination efficiency in unchallenged and cisplatin-treated HAP1-*C1orf112^KO^* and HAP1-*FIGNL1^KO^* cells **(Suppl. Fig 4C)**. We next wondered whether downstream dissolution of HR intermediates by BLM helicase may be responsible for maintaining recombination proficiency in HAP1-*C1orf112^KO^* and HAP1-*FIGNL1^KO^* cells. In sgLUC cells, siRNA-mediated depletion of BLM increased SCE numbers, consistent with the hyper-recombination phenotype observed in Bloom Syndrome (**Suppl. Fig 4D**). Notably, in HAP1-*C1orf112^KO^* and HAP1-*FIGNL1^KO^*, the observed increase in SCEs upon BLM depletion was lost (**Suppl. Fig 4D**). These data suggest that C1orf112 and FIGNL1 only affect recombination when processing of joint molecules by BLM is simultaneously compromised.

In addition to its role in HR, RAD51 is also involved in maintenance of replication forks. DNA fibre assays revealed that replication speed was slightly reduced in unchallenged HAP1-*C1orf112^KO^* cells **(Suppl. Fig 4E)**. Upon treatment with hydroxyurea (HU), HAP1-*C1orf112^KO^* and HAP1-*FIGNL1^KO^* cells displayed fork protection defects to a similar extent as HAP1-*FANCD2^KO^* cells **(Fig. 4E)**, suggesting a role for C1orf112 and FIGNL1 at replication forks. Using iPOND (Dungrawala & Cortez, 2015), C1orf112 and FIGNL1 were shown to be associated to ongoing replication forks in unchallenged conditions **(Fig. 4F)**. C1orf112 and FIGNL1 showed decreased association to replication forks upon fork stalling by HU treatment **(Fig. 4F)**.

To investigate the consequences of C1orf112 or FIGNL1 inactivation on the timely completion of DNA replication, mitotic FANCD2 foci were analysed **(Fig. 4G, Suppl. Fig 4F)**. An increase in mitotic FANCD2 foci was observed in HAP1-*C1orf112^KO^* and HAP1-*FIGNL1^KO^* cells in unchallenged and cisplatin-treated conditions, indicative of transfer replication intermediates into mitosis **(Fig. 4G, Suppl. Fig 4F)**. Moreover, an increase in 53BP1 bodies was observed in interphase cells **(Fig. 4H)**, a sign of unresolved replication stress-induced lesions being transferred into the next cell cycle (Lukas et al., 2011). Overall, these data suggest that inactivation of C1orf112 or FIGNL1 impairs proper replication fork dynamics, potentially through altered RAD51 chromatin dynamics, resulting in mitotic FANCD2 foci and transfer of replication intermediates into mitosis and the next G1-phase.

### Synthetic lethality between ERCC6L and C1orf112 is accompanied by mitotic aberrancies

Because HAP1-*C1orf112^KO^* and HAP1-*FIGNL1^KO^* cells entered mitosis with unresolved replication intermediates **(Fig. 4G)**, we investigated if the synthetic lethality between PICH and C1orf112 could be due to mitotic aberrancies. Doxycycline-inducible shRNAs targeting PICH were introduced into HAP1-*C1orf112^KO^* and HAP1-*FIGNL1^KO^* cells **(Fig. 5A)**. In accordance with the screen, depletion of PICH resulted in a sharp decrease in cell viability of HAP1-*C1orf112^KO^* and HAP1-*FIGNL1^KO^* cells in colony formation assays **(Fig. 5B, Suppl. Fig 5A)**. In line with these findings, flow cytometry analysis of Annexin-V staining, revealed that apoptosis was induced in HAP1-*C1orf112^KO^* and HAP1-*FIGNL1^KO^* cells upon PICH depletion **(Fig. 5C, Suppl. Fig 5B)**. Strikingly, PICH-depleted HAP1-*C1orf112^KO^* and HAP1-*FIGNL1^KO^* cells displayed high levels of DAPI-positive chromatin bridges in anaphase **(Fig. 5D, Suppl. Fig 5C)**. A large fraction of these bridges displayed γH2AX foci, suggesting chromatin bridge breakage **(Fig. 5D, Suppl. Fig 5C)**. The increased numbers of chromatin bridges were accompanied by an increase in 53BP1 bodies in interphase cells **(Fig. 5E, Suppl. Fig 5D)**, and the formation of micronuclei **(Fig. 5F, Suppl. Fig 5E)**. Combined, these results suggest that PICH-depleted HAP1-*C1orf112^KO^* and HAP1-*FIGNL1^KO^* cells undergo aberrant mitosis, resulting in reduced cell viability.

**Figure 5:**
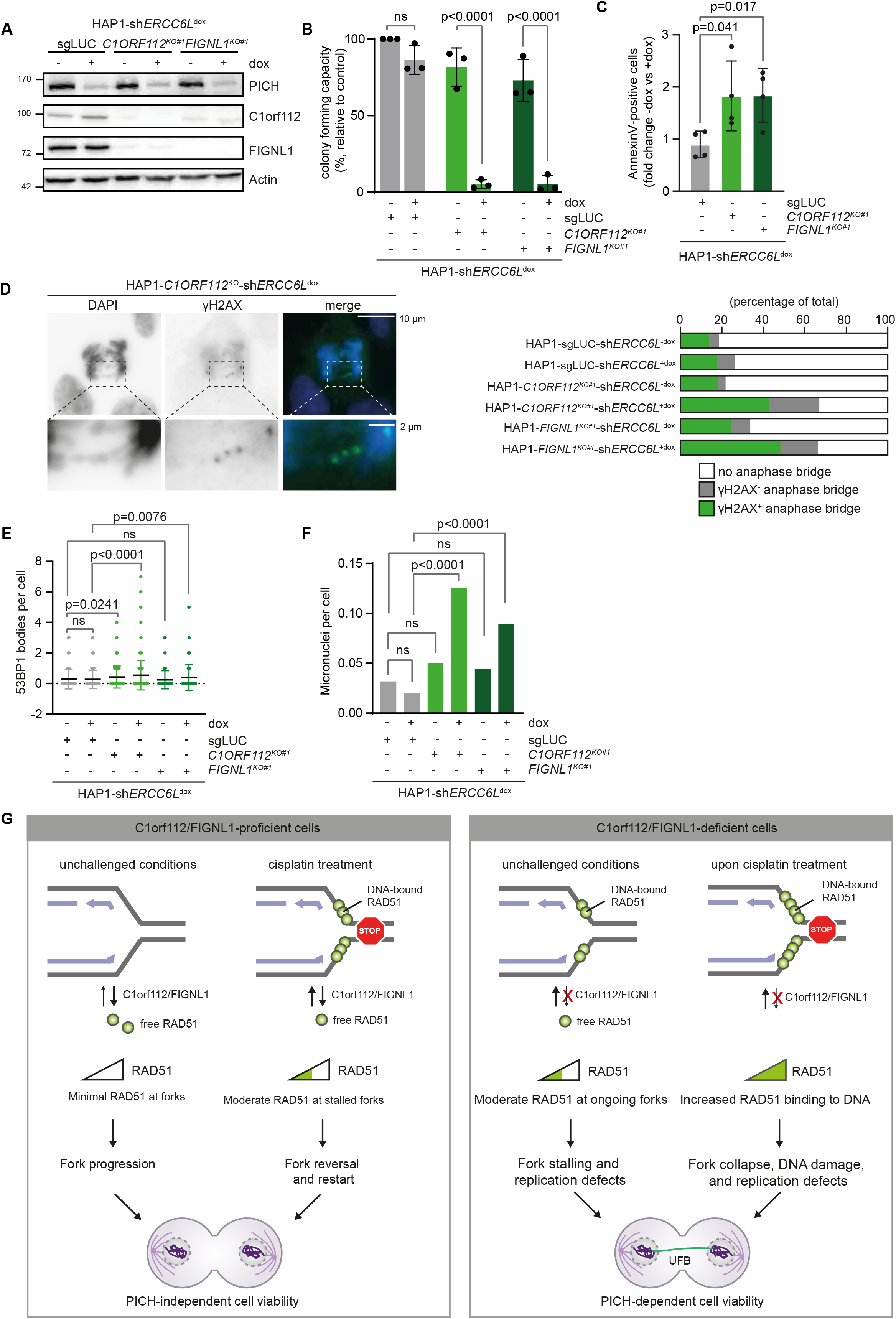
Synthetic lethality between PICH and C1orf112 is associated with mitotic aberrancies. **(A)** shRNAs against *ERCC6L* were introduced in HAP1-sgLUC, HAP1-*C1orf112^KO^* and HAP1-*FIGNL1^KO^* cells, and were induced using 0.1 μM doxycycline for 48 h. Expression of PICH, C1orf112 and FIGNL1 were analysed by western blot. **(B)** HAP1-sgLUC-*shERCC6L*, HAP1-*C1orf112^KO^-shERCC6L* and HAP1-*FIGNL1^KO^-shERCC6L* cells were seeded in 6-well plates, and hairpin expression was induced using 0.1 μM doxycycline. Colonies were fixed and stained after 8 days. Graphs represent means and standard error of the mean (SEM) from three independent replicates. Statistics were performed using two-tailed t-tests. **(C)** HAP1-sgLUC-*shERCC6L*, HAP1-*C1orf112^KO^-shERCC6L* and HAP1-*FIGNL1^KO^-shERCC6L* cells were treated with 0.1 μM doxycycline for 48 h, and apoptosis induction was subsequently measured using Annexin-V/PI flow cytometry staining. Graphs represent means and standard error of the mean (SEM) of four independent replicates. Statistics were performed using two-tailed t-tests. **(D)** HAP1-sgLUC-*shERCC6L*, HAP1-*C1orf112^KO^-shERCC6L* and HAP1-*FIGNL1^KO^-shERCC6L* cells were treated with 0.1 μM doxycycline for 48 h. Mitotic γH2AX foci and mitotic aberrancies were analysed by immunofluorescence staining. At least 30 anaphases were scored per condition. **(E-F)** HAP1-sgLUC-*shERCC6L*, HAP1-*C1orf112^KO^-shERCC6L* and HAP1-*FIGNL1^KO^-shERCC6L* cells were treated with 0.1 μM doxycycline for 48 h. 53BP1 bodies and micronuclei were analysed by immunofluorescence staining. At least 100 cells were scored per condition. **(G)** Model in which C1orf112 and FIGNL1 regulate RAD51 dynamics at replication forks. Inactivation of C1orf112 and FIGNL1 results in impaired RAD51 dynamics, perturbed DNA replication, and consequently UFB formation in mitosis, rendering cells dependent on PICH.

Overall, our data suggest that C1orf112 and FIGNL1 regulate RAD51 dynamics at chromatin, thereby maintaining proper replication fork dynamics **(Fig. 5G)**. In unchallenged conditions, RAD51 should be removed from ongoing replication forks to ensure proper DNA replication. When treated with replication-blocking agents, such as cisplatin, RAD51 is recruited to repair replication intermediates to promote fork reversal and restart **(Fig. 5G)**. In the absence of C1orf112, RAD51 may be increasingly associated with ongoing forks, resulting in moderate replication defects. Upon treatment with cisplatin, RAD51 retention on DNA is further enhanced, possibly preventing proper fork reversal and restart and resolution of repair intermediates or late-stage replication intermediates **(Fig. 5G)**. This model is consistent with a previously established role for FIGNL1 in regulating RAD51 filament dynamics (Matsuzaki et al., 2019). Overall, impaired RAD51 dynamics in *C1orf112* and *FIGNL1*-deficient cells result in replication defects, transfer of replication-derived DNA lesions into mitosis, an increase in UFBs, and a consequently stronger dependence on PICH to maintain genome stability and cell viability **(Fig. 5G)**.

## Discussion

Here we present a genetic context for *ERCC6L* (PICH) essentiality using a genome-wide loss-of-function screen. Genes that displayed synthetic lethality with *ERCC6L* were generally involved in DNA condensation and cohesion (*i.e*. NCAPD3, NCAPG2, NSMCE2) and mitotic resolution of joint DNA molecules (*i.e*. EME1). Some of the identified genes have previously been linked to UFB biology, including NCAPD3, which is part of the condensin II complex and regulates the compaction and disentanglement of sister chromatids (Ono et al., 2003). Mutations in *NCAPD3* result in decatenation failure and UFB formation (Martin et al., 2016). Additionally, EME1 and its interaction partner MUS81 are part of the SLX4-structure-specific tri-nuclease complex, which prevents replication stress-associated UFB formation (Ying et al., 2013). Moreover, the SMC5/6-subunit NSMCE2 has recently been described as a novel player in the rescue of stalled replication forks (Pond et al., 2019). NSMCE2 depletion has been associated with increased numbers of RAD51 foci, mitotic DNA damage and UFB formation (Pond et al., 2019). Finally, the *Schizosaccharomyces pombe* NFATC2IP homolog *Rad60* has been identified as a potential interactor of the SMC5/6 complex, involved in the repair of replication-associated DNA lesions (Boddy et al., 2003; Morishita et al., 2002).

In addition to several previously described genes, we identified synthetic lethality between *ERCC6L* and the uncharacterised open reading frame *C1orf112*, which we identified to interact with FIGNL1. Interestingly, this interaction had previously been described in *Arabidopsis thaliana* (Fernandes et al., 2018), showing that this interaction is conserved from plants to humans. Proteasome-dependent reduction of FIGNL1 protein levels in C1orf112-deficient cells, suggests that C1orf112 has a role in stabilising FIGNL1 protein by protecting it from degradation.

Upon inactivation of either C1orf112 or FIGNL1, a substantial increase in nuclear RAD51 signal in both unchallenged and cisplatin-treated conditions was observed. In humans, RAD51 is the key recombinase involved in the homology search and strand invasion steps of HR (P. Baumann & West, 1998), while RAD51 also binds ssDNA at stalled replication forks to promote fork protection, reversal and restart (Kolinjivadi et al., 2017). Differential RAD51 requirements have been described for HR, fork protection and fork reversal (Kolinjivadi et al., 2017). High levels of RAD51 activity are required to promote BRCA2-mediated strand invasion during HR, whereas fork reversal is BRCA2-independent and requires much lower RAD51 levels (Kolinjivadi et al., 2017). Proper regulation of RAD51 binding to ssDNA is crucial for the preservation of genome stability, as both RAD51 deficiency and RAD51 overexpression result in genome instability (Richardson et al., 2004; Sonoda et al., 1998). We suggest that C1orf112 and FIGNL1 play a role in regulating RAD51 dynamics at replication forks. C1orf112 and FIGNL1 were detected at nascent DNA at unchallenged and stalled replication forks. Moreover, C1orf112-deficiency resulted in defective replication fork progression and protection. We observed prolonged RAD51 retention on DNA upon cisplatin treatment in HAP1-*C1orf112^KO^* and HAP1-*FIGNL1^KO^* cells. Surprisingly, our FRAP data showed increased mobilisation of overall nuclear GFP-RAD51 in *C1orf112^KO^* and *FIGNL1^KO^* cells. Of note, GFP-RAD51 FRAP analysis was conducted on the total pool of nuclear GFP-RAD51 rather than individual RAD51 foci due to technical constraints. Increased mobilisation of RAD51 in response to replication stress has been observed previously, and may represent a separate pool of BRCA2-bound RAD51 that is released upon induction of replication arrest (Yu et al., 2003). Combined, these results suggest that in HAP1-*C1orf112^KO^* and HAP1-*FIGNL1^KO^* cells, nuclear RAD51 is mobilised in response to replication stress, and subsequently remains DNA-bound for prolonged times compared to HAP1 control cells.

A role for FIGNL1 in regulating RAD51 dynamics has been described previously (Matsuzaki et al., 2019; Yuan & Chen, 2013). However, conflicting findings have been reported regarding the requirement of FIGNL1 for DNA recombination (Matsuzaki et al., 2019; Yuan & Chen, 2013). In plants, C1orf112 and FIGNL1 have been described as anti-recombinase proteins, suppressing SCE induction during meiosis (Fernandes et al., 2018; Hu et al., 2017). We find that C1orf112 and FIGNL1 are not required for HR-mediated repair of I-SceI-induced DSBs, in contrast to the RAD51-regulator BRCA2 **(Fig. 4D)** or RAD51 itself (Krajewska et al., 2015). In line with these results, C1orf112 or FIGNL1 are not required for SCE formation *per se*, as spontaneous and cisplatin-induced SCEs still occurred in HAP1-*C1orf112^KO^* cells HAP1-*FIGNL1^KO^* cells. These observations are in agreement with genome-wide studies of HR-mediated DSB repair, in which RAD51 and its upstream regulators (*i.e*. BRCA1, BRCA2, PALB2) are required for DSB repair, but C1orf112 or FIGNL1 are not (Adamson et al., 2012; Słabicki et al., 2010). Joint DNA molecules that arise during HR can either be resolved by the MUS81-EME1 endonuclease complex or ‘dissolved’ by BLM helicase. In BLM-depleted cells, joint molecules need to be resolved, leading to elevated levels of SCEs. Because we observed SCE numbers to be reduced when C1orf112 or FIGNL1 were inactivated in BLM-depleted cells **(Suppl. Fig. 4D)**, these results suggest that loss of C1orf112 or FIGNL1 may negatively affect joint molecule resolution, but not BLM-mediated dissolution, thereby maintaining active HR. Similar observations were made for SLX4-tri-nuclease complex components MUS81 and GEN1, which are not required for HR-mediated repair (Adamson et al., 2012; Słabicki et al., 2010), but are required for SCE formation in BLM-defective cells (Wyatt et al., 2017), and prevent UFB formation (Y. W. Chan et al., 2018). Possibly, aberrant retention of RAD51 on chromatin in *C1orf112^KO^* or *FIGNL1^KO^* cells precludes the recruitment of proteins that mediate joint molecule resolution.

Beyond HR repair, RAD51 is involved in maintaining the stability of replication forks. We find C1orf112 and FIGNL1 to be present at nascent DNA at replication forks under physiological conditions. In this context, changes in RAD51 dynamics due to loss of C1orf112 or FIGNL1 could explain the altered fork progression rates. Impeded fork progression due to inactivation of C1orf112 or FIGNL1, may lead to an accumulation of persistent DNA replication intermediates, observed as FANCD2 foci in mitosis. The inability to resolve joint-molecules during mitosis, either due to perturbed replication or unresolved DNA repair intermediates, results in elevated levels of UFBs and a dependence on PICH. In line with this model, combined inactivation of PICH and C1orf112 or FIGNL1 leads to γH2AX-positive mitotic chromatin bridges, and elevated levels of cell death.

*C1orf112* orthologs have been found in plants, invertebrates and vertebrates, suggesting the gene may have originated in the ancestor of all eukaryotes (Edogbanya et al., 2021). Surprisingly however, no *C1orf112* ortholog has been found in yeast, suggesting another non-related protein may be involved in regulation of yeast RAD51 dynamics. In fact, the yeast Srs2 helicase has been described to perturb RAD51 filament formation, and dissociate RAD51 from the DNA (Antony et al., 2009; Qiu et al., 2013). Notably, no Srs2 ortholog has been found in humans (Marini & Krejci, 2010). Potentially, the C1orf112-FIGNL1 complex plays a similar role in human cells as Srs2 does in yeast, despite showing no sequence homology.

Interestingly, the fork protection defects observed in C1orf112-deficient cells are reminiscent of those observed in RADX-deficient cells (Adolph et al., 2021; Dungrawala et al., 2017). RADX is a ssDNA-binding protein that competes for ssDNA binding at stalled replication forks, and prevents RAD51 filament formation at ongoing replication forks (Adolph et al., 2021; Dungrawala et al., 2017). Loss of RADX results in excessive fork-bound RAD51, and subsequently fork slowing and DBS formation (Adolph et al., 2021; Dungrawala et al., 2017). Despite having a potentially similar function as C1orf112, RADX did not show synthetic lethality with *ERCC6L* in our genetic screen **(Suppl. Data 2)**. This suggests that regulation of RAD51 dynamics alone may not fully account for the observed synthetic lethality between *C1orf112* and *ERCC6L*, and we cannot rule out that C1orf112 has functions beyond FIGNL1 stabilisation and RAD51 filament formation. Indeed, an additional role for C1orf112 in regulating PLK1 activity during early mitosis has been described recently (Xu et al., 2021). In fact, we identified PLK1 as a potential interactor of C1orf112 in our mitotic mass spectrometry data set **(Suppl. Fig. 2D)**.Of note, a role for PICH in the maintenance of replication fork stability during S-phase was recently proposed (Tian et al., 2021), although we did not observe altered replication dynamics in PICH-deficient cells.

Overall, our results show that PICH becomes essential for cell viability in cells that display perturbed replication fork protection, DNA condensation, or mitotic processing of DNA repair intermediates. In particular, we identified a synthetic lethal interaction with *C1orf112*, which in conjunction with FIGNL1, regulates RAD51 dynamics. Cells lacking C1orf112 or FIGNL1 show impaired replication fork progression, an accumulation of DNA replication intermediates and eventually UFB formation.

## Materials and Methods

### Cell lines

HAP1 cells were obtained from Horizon Discovery (Cambridge, UK) and maintained in Iscove’s Modifed Dulbecco’s Medium (IMDM; Gibco), supplemented with 2 mM L-glutamine (Gibco), penicillin/streptomycin (Gibco) and 10% FCS (Gibco). MCF-7 human breast cancer cells were obtained from ATCC and cultured in Roswell Park Memorial Institute medium (RPMI-1640; Gibco) supplemented with penicillin/streptomycin and 10% FCS. Human retinal pigment epithelium (RPE-1) cells and MDA-MB-231 human breast cancer cells were obtained from ATCC and cultured in Dulbecco’s Modified Eagle’s Medium (DMEM; Gibco) with reduced glucose (1 g/L), supplemented with penicillin/streptomycin and 10% FCS. Human embryonic kidney (HEK) 293T cells and HeLa cells were cultured in DMEM medium with 4.5 g/L glucose, supplemented with penicillin/streptomycin and 10% FCS. Cells were maintained at 37°C, 5% CO_2_ and 20% O_2_, and were regularly tested negative for mycoplasma.

### CRISPR/Cas9-mediated gene inactivation

HAP1 *ERCC6L^KO^* cell lines were engineered by Horizon Discovery (*sgERCC6L#1*: GGAGGCATCCCGAAGGTTTC, *sgERCC6L#2*: GGGCTCAAGGCCTCGGCTTC, *sgERCC6L#3*: GGGCTCAAGGCCTCGGCTTC). Single guide RNAs (sgRNAs) against luciferase (*sgLUC*: TGGTGTTCGTTTCCAAAAAG), *C1orf112* (*sgC1orf112#1*: TGGAAATCCAGACCACTCTA, *sgC1ORF112#2*: AAATATTCCTTAGAGTGGTC, *sgC1orf112#3*: ATTTTGGCTTGACAGTCCTC), and *FIGNL1 (sgFIGNL1#1*: TAGGAGCTAGTAGATCCCGA, *sgFIGNL1#2*: GAAGACCCTGATGCACGCTG, *sgFIGNL1#3*: ACCAACCTCAGCGTGCATCA, *sgFIGNL1#4*: CCTATACCCAAGCAAGATGG) were cloned into lentiCRISPRv2 (a gift from Feng Zhang, Addgene plasmid #52961) in a single-step digestion-ligation reaction (2 hours (h) at 37°C, 2 h at 16°C, 4 cycles). sgRNAs were introduced through FuGene-mediated transfections, and HAP1 cells were subsequently selected using 1 μg/mL puromycin for 5 days. HAP1 *FANCD2^KO^* cell lines were generated through insertion of a puromycin cassette into exon 4 of FANCD2 (sgFANCD2: CACCGAGAAGCTCTTTCAGACCCTG), according to a protocol described previously (Lackner et al., 2015).

### RNA interference

Short-hairpin RNAs (shRNAs) targeting *ERCC6L (shERCC6L#1*: GCACTTTAAGACATTGCGAAT, *shERCC6L#2*: TATTCTGAGCACTAGCTTAAT), *C1orf112 (shC1orf112#1*: GCAAGTTTCCTCCAAGCCTTT, *shC1orf112#2*: GCTGCTCACATTTCAGCCTTT), *FIGNL1 (shFIGNL1#1*: AGCCAGGAAACAGATAGTAAT, *shFIGNL1#2*: GCAGAAGAATTACTTCGCAAT), *NCAPD3 (shNCAPD3#1*: GCTCTGTTAGAACTGCCTGAA, *shNCAPD3#2*: CCCATTCAGATAAGCTATAAT), and *NFATC2IP (shNFATC2IP#1*: CCTTATCCCAGATGATCTATC, *shNFATC2IP#2*: ACATTTGCTTGAGGCTTATAC) were cloned into Tet-pLKO-puro or TET-pLKO-neo vectors (gifts from Dmitri Wiederschain, Addgene plasmids #21915 and #21916). Hairpins were introduced by lentiviral transduction using pRSV-Rev (Addgene plasmid # 12253), pMDLg/pRRE (#12251), and pMD2.G (#12259) packaging plasmids (gifts from Didier Trono).

Short interfering RNAs (siRNAs) against *C1orf112 (siC1orf112#1*: GGUCUCAGAAACGACAACCAGGAUA, *siC1orf112#2*: GCCCUGGAGAAUGUUAUCAACUCAU), *FIGNL1 (siFIGNL1#1*: GGAGCCAAAGAUGAUUGAACUUAUU, *siFIGNL1#2*: CAG UCU GGA UUG UCA AUA A), *BRCA2 (siBRCA2#1*: CAU AUU GCA GAA GAG UU ACA UUU GAA, *siBRCA2#2*: GGA ACC AAA UGA UAC UGA UCC AUU A), and *BLM (siBLM*: ACAGGGAAUUCUAUGAAGGAGUUAA) were introduced using oligofectamine transfection reagent (ThermoFisher), according to the manufacturer’s recommendations.

### Haploid genetic screen

HAP1 *sgERCC6L^KO^#2* cells were mutagenized as described previously (Blomen et al., 2015; Heijink et al., 2015). In summary, ~6×10^7^ cells were transduced with pGT gene-trap retrovirus. Mutagenized cells were passaged for 12 days prior to fixation in ‘Fixation Buffer 1’ (Becton Dickinson). Genomic DNA was isolated from ~4×10^7^ cells, and digested using NlaIII and MseI. DNA was subjected to a linear amplification (LAM)-PCR protocol, and sequenced using the Genome Analyzer platform (Illumina). Gene trap insertion mapping and data analysis was performed as described previously (Raaijmakers et al., 2018). In brief, the dataset was normalized to combined insertion counts of four HAP1 wildtype datasets (Blomen et al., 2015). A binomial test was performed for the distribution of sense and antisense orientation insertions in two HAP1 *sgERCC6L^KO^#2* datasets. A gene was considered synthetic lethal with *ERCC6L* when the sense-over-antisense insertion ratio was decreased in both HAP1 *sgERCC6L^KO^#2* datasets compared to the four HAP1 wildtype datasets (Blomen et al., 2015; SRA accession no. SRP058962). To this end, two-sided Fisher’s exact tests were performed for each gene against all wildtype datasets, and the gene was considered as synthetic lethal when it passed all Fisher’s tests (P < 0.05; effect size 20%).

### Depmap analysis and gene set enrichment

CERES-corrected Achilles gene effect scores for *ERCC6L* were obtained from the Cancer Dependency Map (DepMap) portal (Meyers et al., 2017; Tsherniak et al., 2017). Out of 582 cell lines, 96 were identified as PICH-dependent, and 486 as PICH-independent, based on an *ERCC6L* essentiality cut-off score of −0.5 (Dempster et al., 2019; Pacini et al., 2020). Microarray expression data from the Broad Institute Cancer Cell Line Encyclopedia (CCLE) was matched for 71 PICH-dependent and 385 PICH-independent cell lines. T-tests were conducted for 19,624 genes to compare mRNA expression values in PICH-dependent versus PICH-independent cell lines. For each gene, the −log_10_(p-value)·sign(t-statistic) were used for GSEA using GO biological processes database. Positive t-statistic and high −log_10_(p-value) translates to higher mRNA expression in PICH-dependent cells than in PICH-independent cells. In order to compare genome instability scores, CCLE absolute copy number variation data was obtained from the DepMap portal. Standard deviation in genome-wide copy numbers were used as a proxy for genome instability for each cell line.

### MTT Assays

HAP1 cells were seeded in 96-well plates at a concentration of 500 cells/well. Cells were pre-treated with 0.1 μg/mL doxycycline for 4 days where indicated. Cells were treated for 5 consecutive days with the following agents: cisplatin (Accord), mitomycin C (Sigma-Aldrich), hydroxy urea (HU; Sigma-Aldrich), olaparib (Axon Medchem), N-methyl-N’-nitro-N-nitrosoguanidine (MNNG; Sigma-Aldrich), and bleomycin (Pharmachemie). Methyl-thiazol tetrazolium (MTT; Sigma-Aldrich) was added to wells at a concentration of 5 mg/mL for 3 h. Culture medium was removed and formazan crystals were dissolved in DMSO. Absorbance was measured using a Multiskan Sky microplate spectrophotometer (Thermo Fisher Scientific) at a wavelength of 520 nm.

### SRB assays

HAP1 cells were seeded in 96-well plates at a concentration of 400 cells/well. Cells were pre-treated with 0.1 μg/mL doxycycline for 4 days where indicated. Cells were fixed for 16 h at 4°C in 100 μL 10% trichloro acetic acid (TCA; Sigma-Aldrich) 0-7 days after cell seeding. Plates were subsequently washed in water, and air-dried overnight. Cells were then stained for 30 min in 100 μL 0.1% SRB solution (Sigma-Aldrich). Subsequently, plates were washed in 1% acetic acid (Millipore), and SRB dye was solubilised in 200 μL 10 mM Tris Base (pH 10.5). Absorbance was measured using a Multiskan Sky microplate spectrophotometer (Thermo Fisher Scientific) at a wavelength of 520 nm.

### Colony survival assays

HAP1 cells were seeded in 6-well plates at a concentration of 100 cells/well. Cells were pre-treated with 0.1 μg/mL doxycycline for 4 days when indicated. 24 h after seeding, cells were treated with cisplatin (Accord), mitomycin C (Sigma-Aldrich), MNNG (Sigma-Aldrich) or olaparib (Axon Medchem). Cisplatin and mitomycin C were washed away after 24h. After 7-10 days, cells were washed in PBS and fixed for 20 min in a 0.2% Coomassie Brilliant Blue (Bio-Rad) solution containing 50% methanol (Merck) and 14% acetic acid (Merck). Plates were washed in water, and air-dried overnight.

### TIDE survival assays

Doxycycline-inducible shRNAs targeting *C1orf112, NFATC2IP* or *NCAPD3* were introduced into HAP1 *ERCC6L^WT^* and HAP1 *ERCC6L^KO^#2* cells. HAP1 *ERCC6L^WT^* and HAP1 *ERCC6L^KO^#2* cells were seeded in a 1:1 ratio in medium supplemented with 0.1 μg/mL doxycycline. After 14 days, genomic DNA was isolated using PureLink Genomic DNA Mini Kit (Invitrogen). TIDE sequencing of the *ERCC6L* locus (*ERCC6L*-FW: AGGGATCACAGACTGAGGCAGCAC, *ERCC6L*-RV: CTCAAATGTCTCCTCTTGCCACGCC) was performed as described previously (Brinkman et al., 2014).

### Western blot

Cells were lysed in Mammalian Protein Extraction Reagent (MPER; Thermo Scientific), supplemented with Halt protease inhibitor and Halt phosphatase inhibitor (Thermo Scientific). Protein concentrations were measured using Pierce BCA Protein Assay Kit (Thermo Scientific). Proteins were separated by SDS-PAGE gel electrophoresis, transferred to PVDF membranes (Immobilon), and blocked in 5% skimmed milk (Sigma-Aldrich) in 0.05% TBS-Tween (Sigma-Aldrich). Immunodetection was performed with antibodies directed against PICH (Novus Biologicals, NBP2-13969, 1:500), C1orf112 (Atlas Antibodies, HPA023778, 1:250), FIGNL1 (Atlas Antibodies, HPA055542, 1:250), NCAPD3 (Bethyl, A300-604A, 1:500), NFATC2IP (Abcam, ab169140, 1:500), RAD51 (GeneTex, gtx70230, 1:500), BRCA2 (Calbiochem, OP95, 1:500), FANCD2 (Novus Biologicals, NB100-182, 1:500), γH2AX (Cell Signaling, 9718, 1:1000), phospho-KAP1 (Bethyl, A300-767A, 1:1000), PCNA (Santa Cruz, sc-56, 1:1000), Lamin B1 (Cell Signaling, 68591, 1:1000), Histone H3 (Abcam, ab176840, 1:2000), HSP90 (Santa Cruz, sc-13119, 1:1000), β-Actin (MP Biomedicals, 69100, 1:1000), and Vinculin (Abcam, ab129002, 1:1000). Horseradish peroxidase (HRP)-conjugated secondary antibodies (DAKO, 1:2000) were used for visualization using chemiluminescence (Lumi-Light, Roche Diagnostics) on a Bio-Rad bioluminescence device, equipped with Quantity One/ChemiDoc XRS software (Bio-Rad). When indicated, cells were treated with 10 μM proteasome inhibitor MG-132 (Sigma-Aldrich) 3h prior to harvest. Cytosolic, nuclear and chromatin-bound subcellular fractions were obtained using Subcellular Protein Fractionation Kit (Thermo Scientific), according to the manufacturer’s recommendations.

### Flow cytometry

HAP1 cells were seeded in 6-well plates at a density of approximately 2·10^5^ cells/well, and treated with 1 μM cisplatin (Accord) for 24 h when indicated. For cell cycle analyses, cells were fixed in icecold 70% ethanol, washed in 0.05% PBS-Tween (Sigma-Aldrich), and incubated with primary antibodies against phosphor-MPM2 (Millipore, 05-368) overnight at 4°C. Alexa Fluor 488-conjugated secondary antibody incubation was performed for 1 h at room temperature. Cells were incubated with 50 μg/mL propidium iodide (PI; Thermo Scientific) and 100 μg /mL RNAse A (Sigma-Aldrich) in 0.05% PBS-Tween for 10 minutes (min). S-phase populations were identified by incorporation of 10 μM 5-ethynyl-2’-deoxyuridine (EdU; Sigma-Aldrich) 45 min before fixation. Click-iT reaction cocktail with Alexa Fluor 647-conjugated azide was prepared according to the manufacturer’s recommendations (Invitrogen). For detection of apoptotic cells, cells were harvested in Annexin V binding buffer (BD Pharmingen) after 48 h of doxycycline treatment, and incubated with Alexa Fluor 488-conjugated Annexin V (Invitrogen) and 50 μg/mL PI for 15 min. At least 10,000 events per sample were analysed on an Agilent Quanteon (Agilent Technologies). Data analysis was performed in FlowJo software (Becton Dickinson).

### Immunoprecipitation and mass spectrometry

HEK293T cells were grown in 10-cm culture dishes, and transfected with 5 μg pBABE-hygro-FLAG-C1orf112 or pBABE-hygro-FLAG-FIGNL1 plasmid using calcium phosphate precipitation (Kingston et al., 2003). When indicated, cells were treated with 0.1% DMSO (Sigma-Aldrich), 0.2 μM aphidocolin (Sigma-Aldrich) for 24 h, 1 μM cisplatin (Accord) for 24 h, or 200 ng/mL nocodazole (Sigma-Aldrich) for 16 h. Cells were collected in M-PER Mammalian Protein Extraction Reagent (Thermo Scientific) supplemented with Halt Protease Inhibitor Cocktail (Thermo Scientific) and Halt Phosphatase Inhibitor Cocktail (Thermo Scientific). For each condition, 1 mg of protein lysate was incubated overnight with 30 μL Anti-FLAG M2 Magnetic Beads (Sigma-Aldrich) at 4°C. Beads were washed three times in TBS according to the manufacturer’s recommendations. Proteins were eluted in 25μL 2x Laemmli Sample Buffer without β-mercaptoethanol for 5 min at 96°C. Samples were loaded onto an SDS-PAGE gel, and a single fraction was isolated for in-gel protein digestion as described previously (Shevchenko et al., 2007). Peptides were analysed using an Ultimate 3000 HPLC system (Thermo Fisher Scientific) coupled to a Q-Exactive-Plus mass spectrometer with a NanoFlex source (Thermo Fisher Scientific), equipped with a stainless-steel emitter. Peptides were mapped in MaxQuant, and analysis was performed in Perseus software.

### Isolation of proteins on nascent DNA (iPOND)

Isolation of proteins on nascent DNA (iPOND) was performed as described previously (Dungrawala & Cortez, 2015). HAP1 cells were treated with 10 μM EdU for 10 min. For chase conditions, cells were treated with 10 μM thymidine for 1 h following EdU treatment. Fork stalling was induced by treatment with 4 mM HU for 2 h. Cells were cross-linked in formaldehyde for 20 min, quenched in 1.25 M glycine, and subsequently harvested from 15-cm dishes. 1·10^8^ cells were permeabilised in 0.25% Triton-X100 for 30 min, and washed in 0.5% BSA. Click-iT reaction cocktail with biotin-conjugated azide was prepared according to the manufacturer’s recommendations (Invitrogen), and cells were incubated for 80 min. Cells were lysed in SDS-lysis buffer, and sonicated for 15 min. Capture of biotin-tagged nascent DNA was performed by incubating samples with streptavidin sepharose high performance (GE Healthcare Life Sciences) agarose beads for 16 h. Beads were washed in lysis buffer and 1 M NaCl, and proteins were eluted in 2x Laemmli sample buffer at 95°C for 25 min.

### Immunofluorescence microscopy

HAP1 cells were seeded on coverslips at a density of 2·10^5^ cells/well, and were allowed to attach for 24 h. When indicated, cells were subsequently treated with 1 μM cisplatin (Accord) for 24 h, 10 μM EdU (Sigma-Aldrich) for 45 min, or 5μM CDK inhibitor RO-3306 (Axon Medchem) for 4 h. Cells were fixed in 2% paraformaldehyde (PFA; Sigma-Aldrich), permeabilised in 0.1% Triton X-100 (Sigma-Aldrich), and blocked in 3% BSA (Serva). Coverslips were incubated overnight at 4°C with primary antibody against PICH (Novus Biologicals, NBP2-13969, 1:500), γH2AX (Cell Signaling, 9718, 1:1000), RAD51 (GeneTex, gtx70230, 1:500), FANCD2 (Novusbio, NB100-182, 1:200) and 53BP1 (Novusbio, NBP2-25028 1:200). Cells were then incubated with Alexa Fluor 488 or Alexa Fluor 647-conjugated secondary antibodies for 1 h at room temperature and mounted with Vectashield antifade mounting medium with DAPI (Vectorlabs). For RAD51 detection, 2% PFA was supplemented with 0.1% Triton X-100, and permeabilization was performed in 0.5% Triton X-100. For FANCD2 detection, cells were pre-extracted for 1 min in PEM buffer (100 mM PIPES pH 6.9, 10 mM EGTA, 1 mM MgCl_2_, 0.2% Triton-X100). For EdU detection, permeabilization was performed in 0.5% Triton, Click-iT reaction cocktail with Alexa Fluor 647-conjugated azide was prepared according to the manufacturer’s recommendations (Invitrogen). Slides were imaged using Leica DM4000B fluorescence microscope or Leica TCS SP2 confocal microscope.

### Fluorescence recovery after photobleaching (FRAP)

HAP1 WT and HAP1 *C1orf112^KO^*#1 cells were seeded in glass-bottom 6-cm dishes, and transfected with 5 μg GFP-RAD51 plasmid (kindly provided by Prof. R. Kanaar, Erasmus University, Rotterdam)(Srivastava et al., 2009) using FuGENE HD Transfection Reagent, according to manufacturer’s recommendations (Promega). 48 h after transfection, cells were treated with 100 μM cisplatin for 4 h when indicated. FRAP analysis was performed with a Zeiss LSM 780 NLO AxioObserver.Z1 inverted microscope coupled to a Chameleon Vision compact OPO two photon laser. Square regions of interest with a fixed height of 28 pixels were drawn along the shortest axis of the nucleus. 100 pre-bleach frames were acquired, followed by a 1 sec bleach pulse with the 488 laser at 100% laser power. Fluorescence recovery within the region of interest was measured every 26 ms for 900 frames. Recovery data was normalised to pre-bleach fluorescence levels, and the mean intensity of at least 20 individual cells per condition were plotted, as described previously (Houtsmuller et al., 1999). Cells with exorbitantly high (> 60 AU) or low (< 20 AU) pre-bleach fluorescence levels were manually excluded from the analysis.

### Reverse transcription quantitative real-time PCR (RT-qPCR)

RNA was isolated from HeLa and RPE-1 cells using RNeasy mini kit, according to the manufacturer’s recommendations (Qiagen). *C1orf112* expression levels were determined using SYBR Green RT-PCR Master Mix and RT Mix, according to manufacturer’s recommendations (QuantiTect). Primers were designed to amplify transcript ENST00000286031 (FW: GCCTTTCATGGTTTGTTCTGAG, RV: TTGAATATGCTCAATCCAACTGTC, amplifies ENST00000413811 as a side product), ENST00000359326 (FW: TATGTCTCAGGAAGGCTGGAGTC, RV: TGAGACCAATCAAGAATGTGCTGC), ENST00000459772 (FW: CCTCTCATTGTCTCAACATTCTG. RV: TAAGAGCTGCACCATTGTTTGTA, amplifies ENST00000496973 and ENST00000466580 as side products), ENST00000496973 (FW: CCTCTCATTGTCTCAACATTCTG, RV: ACAGTCTTTGGTAACACTTGTG), ENST00000472795 (FW: GCACCAGTTTGTTCTGAGTAATG, RV: TTGAATATGCTCAATCCAACTGTC), ENST00000413811 (FW: GGAAGTTTATAATTAAGATAATGCTGAC, RV: CTCTGATTCTTGCTGTACTTTG), and ENST00000498289 (FW: TATCGCCAAGCAACCATGTG, RV: TAGAACACGTTTCCTCTGTCTCC). A primer pair amplifying all transcript variants (FW: TTACAAACAATGGTGCAGCTC, RV: CACAGATTCACTATTCTTACAATGC) was included as a positive control. All reactions were normalised to GAPDH house keeping control reactions (FW: CACCACCACGGAGAACGCTGG, RV: CCAAAGTTGTCATGGATGACC).

### HR assays

HeLa pDR-GFP cells were seeded in 6-well plates, and transfected with 133 nM siRNA targeting *BRCA2, C1orf112* or *FIGNL1* (see sequences above) using Oligofectamine (Thermo Scientific) according to the manufacturer’s recommendations. After 24 h, cells were transfected with pCAGGS-3NLS-I-SceI plasmid to induce a DNA break in the GFP locus, as described previously (Krajewska et al., 2015). Cells were analysed by flow cytometry 48 h after I-Sce1 transfection, using LSR-II flow cytometer (Becton Dickinson, Franklin Lakes, NJ, USA). Three biological replicates were included for each siRNA, statistical analysis was performed using unpaired, two-sided student t-tests.

### Sister chromatid exchange assays

HAP1 cells were treated for 48 h with 10 μM BrdU. When indicated, cells were transfected with siRNA against *BLM* at 24 h prior to BrdU treatment. Alternatively, cells were treated with 1 μM cisplatin simultaneously with BrdU treatment. Colcemid was added during the last 3 h of BrdU treatment at a concentration of 100 ng/mL. Cells were inflated in a hypotonic 0.075 M KCl solution, and subsequently fixed in 3:1 methanol:acetic acid solution. Metaphase spreads were made by dripping the cell suspensions onto microscope glasses from a height of approximately 30 cm. Slides were stained with 10 μg/mL bis-Benzimide H 33258 (Sigma) for 30 minutes, exposed to 245 nM UV light for 30 min, incubated in 2x SSC buffer (Sigma) at 60°C for 1 h, and stained in 5% Giemsa (Sigma) for 15 min.

### DNA fibre assays

HAP1 wildtype, HAP1 *C1ORF112^KO^* and HAP1 *FIGNL1^KO^* cells were seeded in 6-well plates at a concentration of 4·10^5^ cells/well, and incubated for 24 h. For fork progression assays, cells were pulse-labelled with 25 μM 5-iodo-2’-deoxyuridine (IdU; Sigma-Aldrich) for 20 min, washed thrice in full media, and subsequently pulse-labelled with 250 μM 5-chloro-2’-deoxyuridine (CldU; Sigma-Aldrich) for 20 min. For fork protection assays, cells were pulse-labelled with 25 μM IdU for 20 min, washed three times, and subsequently incubated with 250 μM CldU in the presence of 4 mM HU (Sigma-Aldrich) for 3 h. Cells were harvested by trypsinisation and diluted to a concentration of 5·10^5^ cells/mL. Cells were subsequently lysed in lysis buffer (0.5% sodium dodecyl sulfate, 200 mM Tris pH 7.4, 50 mM EDTA) and allowed to spread by gravity flow on tilted microscopy slides. Slides were air-dried, subsequently fixed in methanol/acetic acid (3:1) for 15 min, and denatured for 1.5 h in 2.5 M HCl. IdU was detected using mouse anti-BrdU (BD Biosciences, 347580, 1:250), CldU was detected using rat anti-BrdU (Abcam, ab6326, 1:1000). Slides were incubated with primary antibodies for 1-2 h at room temperature, followed by incubation with Alexa Fluor 488- or 647-conjugated secondary antibodies (1:500) for 1.5 h. Images were acquired on a Leica DM-6000RXA fluorescence microscope, equipped with Leica Application Suite software. The lengths of CIdU and IdU tracks were measured using ImageJ software. Two-sided Mann-Whitney tests were used for statistical analysis.

### Single cell whole genome sequencing

HAP1 wildtype, HAP1 *C1ORF112^KO^* and HAP1 *FIGNL1^KO^* cells were treated with 1 μM cisplatin for 96 h. Subsequently, cells were incubated in mild lysis buffer, and single G_1_ nuclei were sorted into 96□well plates, using a Hoechst/propidium iodide double staining. Illumina□based library preparation was performed as described previously (van den Bos et al., 2016), using a Bravo automated liquid handling platform (Agilent Technologies, Santa Clara, CA, USA). Single□cell libraries were pooled and sequenced on an Illumina NextSeq 500 sequencer (Illumina, San Diego, CA, USA). Sequencing data were analysed using Aneufinder software (Bakker et al., 2016). Modal copy number states of cells were determined per sample and per bin, and bins that deviated from the modal copy number state were identified. The genomic instability scores were assessed per cell, by determining the fraction of bins that deviate from the modal copy number for that sample. All sequencing data have been deposited at the European Nucleotide Archive under accession PRJEB54793.

## Data availability

The mass spectrometry proteomics data have been deposited to the ProteomeXchange Consortium via the PRIDE partner repository with the dataset identifier PXD035876. Strand-seq data are available via the European Nucleotide Archive (ENA) with identifier RJEB54793.

## Acknowledgments

This work was supported by a grant from the European Research Council (ERC-Consolidator grant #681572 ‘TENSION’) to M.A.T.M.v.V. We thank Thijn Brummelkamp and Maarten Hekkelmans (Netherlands Cancer Institute, Amsterdam) for technical support and Roland Kanaar (ErasmusMC, Rotterdam) for sharing reagents. We thank members of the Medical Oncology Department for discussions and feedback on the manuscript.

---

**Suppl. Figure 1: *ERCC6L* is not essential for cell viability in a majority of tumour cell lines. (A)** HAP1-*ERCC6L^WT^* and HAP-*ERCC6L^KO^* cells were seeded in 6-well plates, and colony formation was measured after 7 days by Coomassie Brilliant Blue staining. Pictures represent a single representative well out of three replicates. **(B)** Cell cycle profiles of HAP1-*ERCC6L^WT^* and HAP1-*ERCC6L^KO^* cells were generated by flow cytometry analysis of EdU, MPM2 and PI signal. Representative dot plots of EdU, MPM2 and PI flow cytometry staining are presented. Percentage of mitotic cells was measured by analysis of MPM2-positive cells. Bar graphs represent mean and standard error of the mean (SEM) of three replicates. **(C)** Hairpins against *ERCC6L* were introduced into MDA-MB-231 and MCF-7 human breast cancer cell lines, and induced by treatment with 0.1 μg/mL doxycycline (dox) for 96 h. PICH protein expression levels were analysed in whole-cell lysates by western blot. **(D)** Cell growth of MDA-MB-231-*shERCC6L* and MCF-7-*shERCC6L* cells was monitored for 7 days using SRB staining. Data points represent the mean and standard error of the mean (SEM) of three replicates. **(E)** Cell viability of MDA-MB-231-*shERCC6L* and *MCF-7-shERCC6L* cells was monitored for 7 days by measuring MTT conversion. Data points represent the mean and standard error of the mean (SEM) of three replicates. **(F)** Deviations from the modal copy number were analysed for 21 HAP1-*ERCC6L^WT^* and 11 HAP1-*ERCC6L^KO^* single-cell sequencing libraries. **(G)** *ERCC6L* CERES-corrected gene essentiality scores for 582 cell lines from Depmap. Cell lines with CERES scores below −0.5 were considered PICH-dependent. Cell lines were categorized by their tissue of origin. **(H)** *ERCC6L* mRNA expression levels from 71 PICH-dependent and 385 PICH-independent cell lines from the CCLE data. Statistics were performed using a two-tailed Mann-Whitney test. **(I)** Genome instability scores were calculated for 71 PICH-dependent and 385 PICH-independent cell lines. Statistics were performed using a two-tailed Mann-Whitney test. **(J)** mRNA expression levels of 19,624 genes were compared in 71 PICH-dependent and 385 PICH-independent cell lines using t-tests, and gene set enrichment analysis was performed. Z-transformed p-values are plotted on the x-axis, larger triangle size indicates higher significance. **(K)** Hairpins targeting *C1orf112, NCAPD3* and *NFATC2IP* were introduced into HAP1-*ERCC6L^WT^* and HAP1-*ERCC6L^KO^* cells, and induced by treatment with 0.1 μg/mL doxycycline (dox) for 48 h. C1orf112, NCAPD3, and NFATC2IP protein expression levels were analysed in whole-cell lysates by western blot. **(L)** HAP1-*ERCC6L^WT^* and HAP1-*ERCC6L^KO^* cells expressing hairpins targeting *C1orf112, NCAPD3* and *NFATC2IP* were mixed in a 1:1 ratio, and abundance of the *ERCC6L* mutation was monitored over time by TIDE analysis. Graphs represent abundance of HAP1-*ERCC6L^KO^* cells after 14 days of co-culture in a single representative replicate.

**Suppl. Figure 2: The largest protein-coding isoform of C1orf112 interacts with FIGNL1. (A)** Nine transcript variants have been described for C1orf112, including five protein-coding isoforms. Exon distribution is plotted as red squares. **(B)** mRNA expression levels of the C1orf112 transcripts were determined using ENCODE GTEx data. TPM data in each sample was normalised to total C1orf112 expression in that sample. **(C)** expression levels of the various C1orf112 transcript variants were determined by one-step RT-qPCR analysis of HeLa and RPE-1 cells. Graphs represent the mean of three technical replicates. **(D)** HEK293T cells were transiently transfected with FLAG-C1orf112 or FLAG-FIGNL1 construct, and treated with DMSO, 0.2 μM APH for 24 h, 1 μM cisplatin for 24 h, or 200 ng/mL nocodazole for 16 h. Immunoprecipitation was performed in whole-cell lysates using anti-FLAG M2 beads. Proteins were digested in-gel, and analysed by HPLC-MS. log_2_(LFQ Intensity) scores are plotted for C1orf112, FIGNL1 and PLK1. log_2_(LFQ Intensity) scores in DMSO-treated cells represent the mean of four independent datasets, in APH-treated cells the mean of three datasets, and in cisplatin-treated cells two independent experiments. **(E)** HEK293T cells were transfected with FLAG-FIGNL1 construct, and IP reactions were performed in whole-cell lysates using anti-FLAG M2 beads. C1orf112 and FIGNL1 levels in input and IP fraction were analysed by western blot. **(F)** Percentage of sense insertions in the FIGNL1 locus. Datapoints represent four independent HAP1-*ERCC6L^WT^* datasets and two independent HAP1-*ERCC6L^KO^* datasets. **(G)** HAP1-*C1orf112^KO^* and HAP1-*FIGNL1^KO^* cells were treated with 10 μM MG-132 proteasome inhibitor for 3 h. C1orf112 and FIGNL1 protein expression levels were analysed in whole-cell lysates by western blot, and quantified in ImageJ. Datapoints represent three independent replicates, statistics were performed using t-tests. **(H)** Function predictions for C1orf112 and FIGNL1 were performed using permutated Z-scores from GenetICA. Correlation was expressed as Spearman’s r score. **(I)** CERES-corrected gene essentiality scores for C1orf112 and FIGNL1 were obtained from the Depmap consortium from 582 cell lines. Correlation was expressed as Spearman’s r score.

**Suppl. Figure 3: Sensitivity profiles to DNA damaging agents for C1orf112, FIGNL1 and FANCD2-deficient cells. (A)** HAP1-*C1orf112^KO^*, HAP1-*FIGNL1^KO^* and HAP1-*FANCD2^KO^* cells were treated with MNNG for 7 days, and colonies were analysed by Coomassie Brilliant Blue staining. Graphs represent mean and standard error of the mean (SEM) of three independent biological replicates. **(B)** HAP1-*C1orf112^KO^*, HAP1-*FIGNL1^KO^* and HAP1-*FANCD2^KO^* cells were treatmed with cisplatin, olaparib, mitomycin C, hydroxy urea, MNNG and bleomycin at 24 h after seeding. Cell viability was measured by MTT conversion after 5 days. Data points represent the mean and standard error of the mean (SEM) of three replicates. **(C)** HAP1-*C1orf112^KO^* and HAP1-*FIGNL1^KO^* cells were treated with 1 μM cisplatin for 24 h, and FANCD2 ubiquitination levels were analysed by western blot. **(D)** HAP1-*C1orf112^KO^*, HAP1-*FIGNL1^KO^* and HAP1-*FANCD2^KO^* cells were treated with 1 μM cisplatin for 24 h, and whole cell lysates were harvested at 0 h, 8 h and 24 h after washout of cisplatin. FANCD2, RAD51, and phospho-KAP1 levels were analysed by western blot. **(E)** Copy number alterations (CNAs) were analysed using single-cell sequencing of 28 HAP1, 19 HAP1-*C1orf112^KO^* and 28 HAP1-*FIGNL1^KO^* libraries. If indicated, cells were treated with 1 μM cisplatin for 96 h. CNA analysis in cisplatin-treated cells was performed using 24 HAP1, 31 HAP1-*C1orf112^KO^* and 27 HAP1-*FIGNL1^KO^* libraries. CNAs were defined as deviations from the modal copy number.

**Suppl. Figure 4: C1orf112 and FIGNL1-deficient cells display altered RAD51 dynamics. (A)** HAP1-*C1orf112^KO^*, HAP1-*FIGNL1^KO^* and HAP1-*FANCD2^KO^* cells were treated with 1 μM cisplatin for 24 h, and cells were fixed at 0 h, 8 h and 24 h after washout of cisplatin. RAD51 nuclear intensity was analyzed using ImageJ. At least 100 cells per condition were scored. Statistics were performed using two-tailed Mann-Whitney tests. **(B)** HAP1 ctrl and HAP1-*C1orf112^KO^* cells were transfected with GFP-RAD51 plasmid. 24 h after transfection, cells were treated with 100 μM cisplatin for 4 h. After GFP bleaching, fluorescence recovery was measured for 900 frames. Graphs represents the mean recovery of at least 20 individual nuclei per condition. Data was normalised to initial pre-bleach fluorescence. **(C-D)** HAP1-sgLUC, HAP1-*C1orf112^KO^* and HAP1-*FIGNL1^KO^* cells were treated with BrdU for 48 h. When indicated, cells were treated with 1 μM cisplatin for 48 h or transfected with siRNA against BLM 48 h prior to fixation. Cells were synchronised with colcemid, and metaphase spreads were analysed using Giemsa stain. At least 30 metaphases were scored per condition. Statistics were performed using two-tailed Mann-Whitney tests. **(E)** HAP1-sgLUC, HAP1-*C1ORF112^KO^*, HAP1-*FIGNL1^KO^* and HAP1-*FANCD2^KO^* cells were subsequently pulse-labelled with IdU for 20 min and CldU for 20 min. Lengths of at least 200 DNA fibres were measured per condition. Statistics were performed by two-tailed Mann-Whitney tests. **(F)** HAP1-sgLUC, HAP1-*C1ORF112^KO^* and HAP1-*FIGNL1^KO^* cells were treated with 1 μM cisplatin for 24 h, and subsequently FANCD2 foci were analysed by immunofluorescence staining. Representative images are shown for FANCD2 foci.

**Suppl. Figure 5: Synthetic lethality between PICH and C1orf112 is associated with mitotic aberrancies. (A)** HAP1-sgLUC-*shERCC6L*, HAP1-*C1orf112^KO^-shERCC6L* and HAP1-*FIGNL1^KO^-shERCC6L* were treated with 0.1 μM doxycycline. Colony formation was measured after 8 days. Pictures represent representative wells out of three replicates. **(B)** HAP1-sgLUC-*shERCC6L*, HAP1-*C1orf112^KO^-shERCC6L* and HAP1-*FIGNL1^KO^-shERCC6L* cells were treated with 0.1 μM doxycycline for 48 h, and apoptosis induction was subsequently measured using Annexin-V/PI flow cytometry staining. Representative dot plots are shown. **(C-E)** HAP1-*shERCC6L*, HAP1-*C1orf112^KO^-shERCC6L* and HAP1-*FIGNL1^KO^-shERCC6L* cells were treated with 0.1 μM doxycycline for 48 h. Mitotic γH2AX foci, mitotic aberrancies, 53BP1 bodies and micronuclei were analysed by immunofluorescence staining.

**Suppl. Data 1: Geneset enrichment analysis of PICH-dependent and PICH-independent cell lines**

**Suppl. Data 2: Genetrap mapping in WT and *ERCC6L-mutant* HAP1 cell lines**

**Suppl. Data 3: GenetICA gene function predictions**

**Suppl. Data 4: Peptides from C1orf112 IP-MS analysis**

## Notes

### Competing Interest Statement

The authors have declared no competing interest.

