## Supplementary figures and images for "The FIGNL1-interacting protein C1orf112 is synthetic lethal with PICH and mediates RAD51 retention on chromatin"

### Figure S1

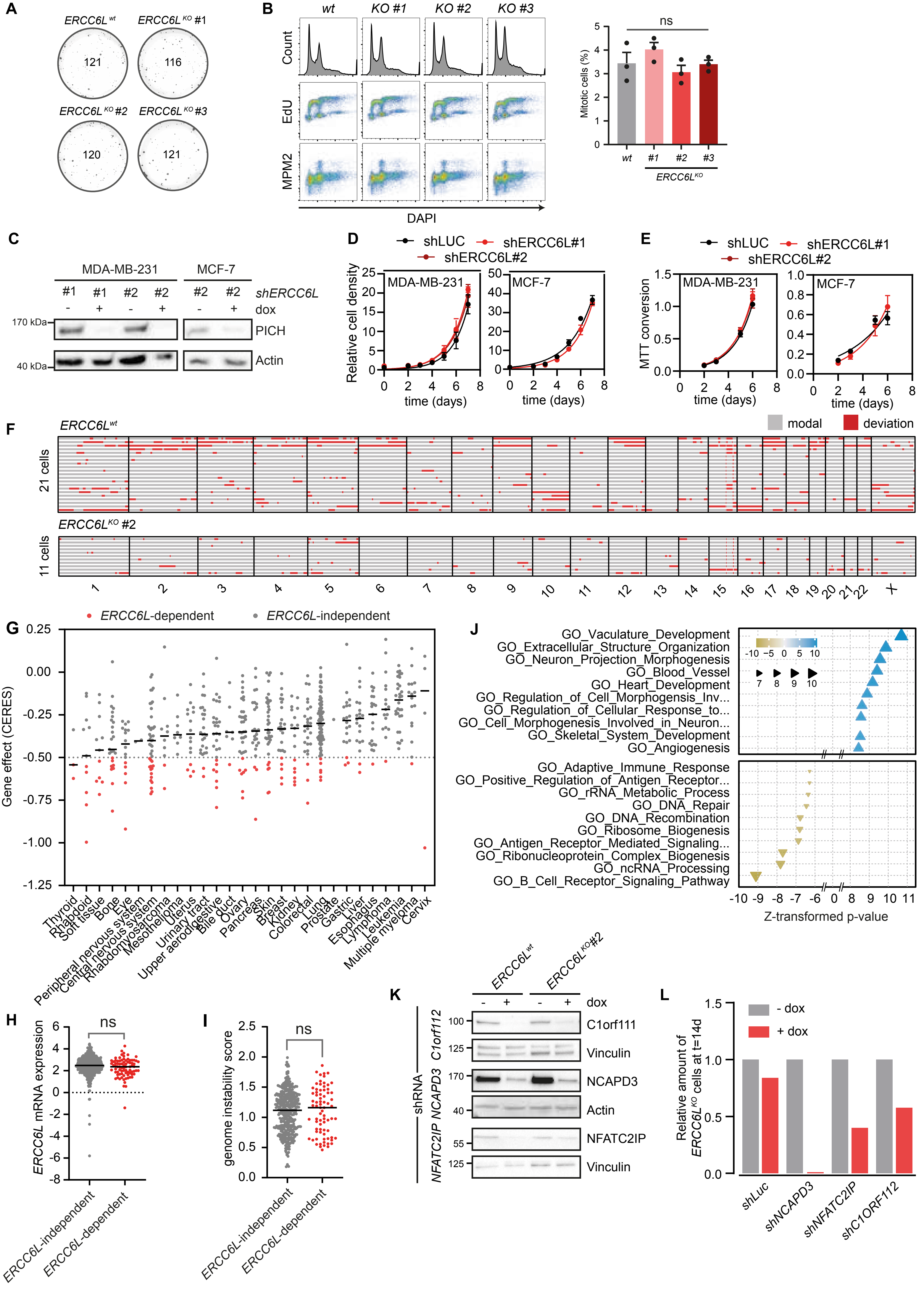

### Figure S2

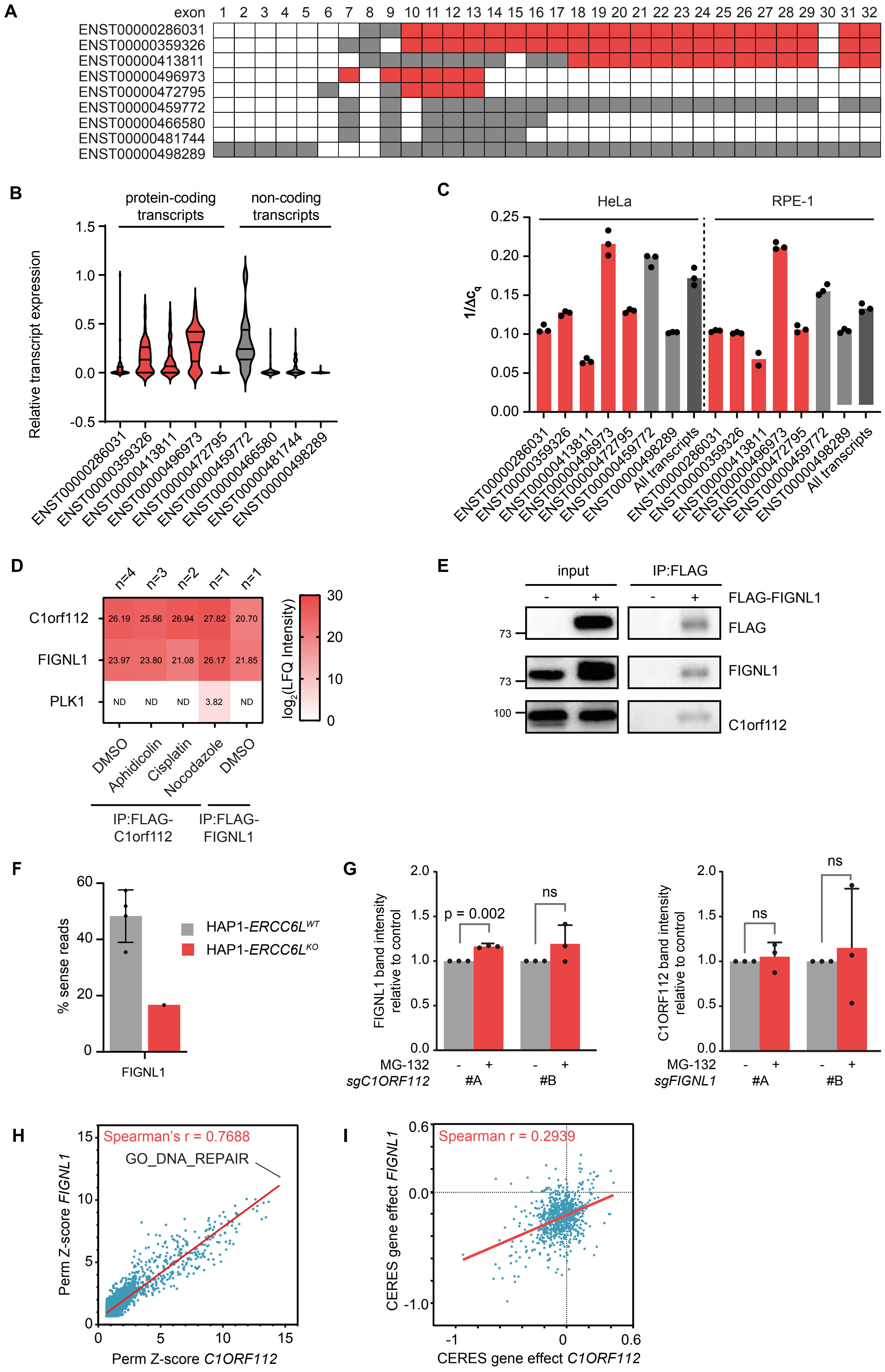

### Figure S3

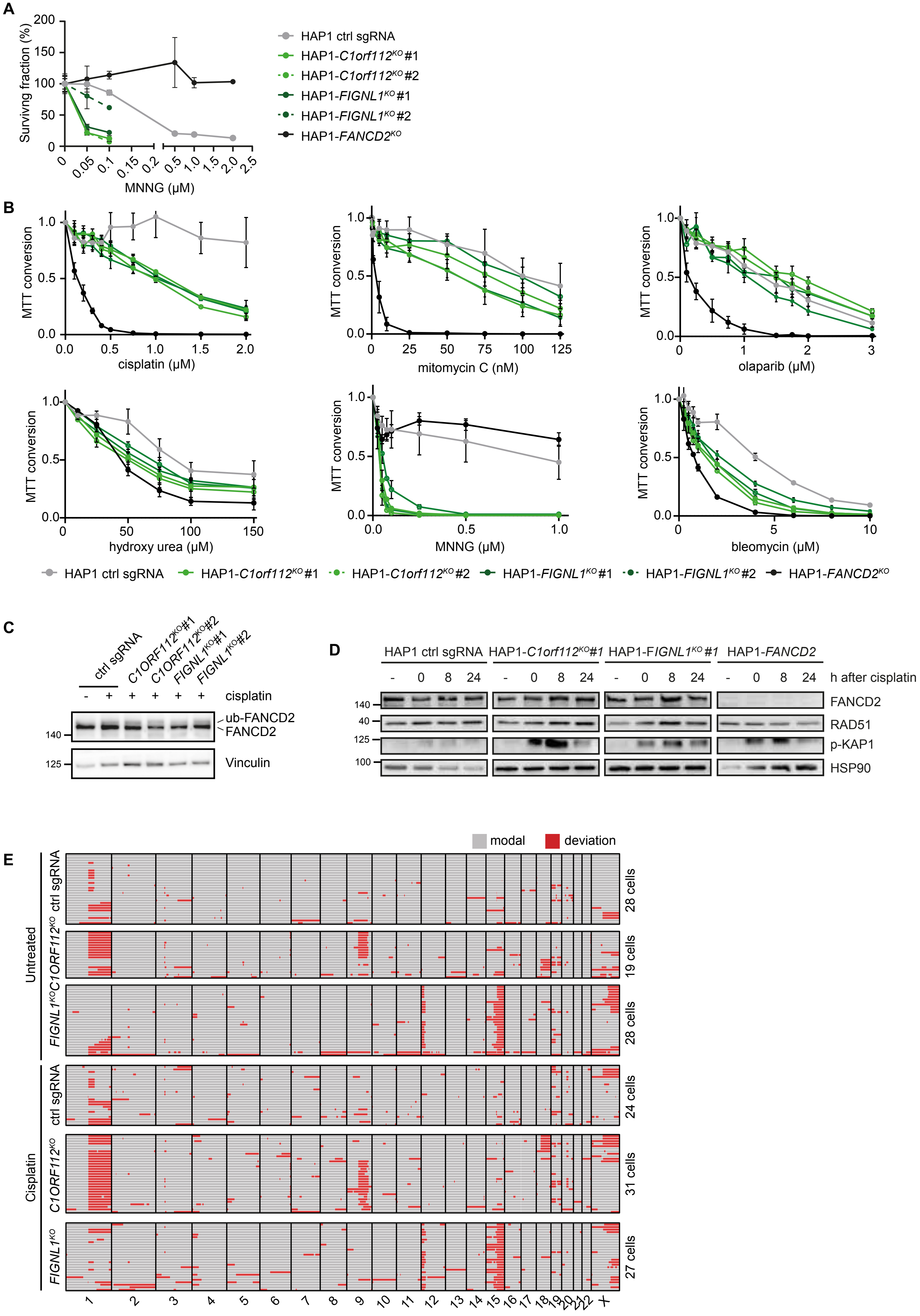

### Figure S4

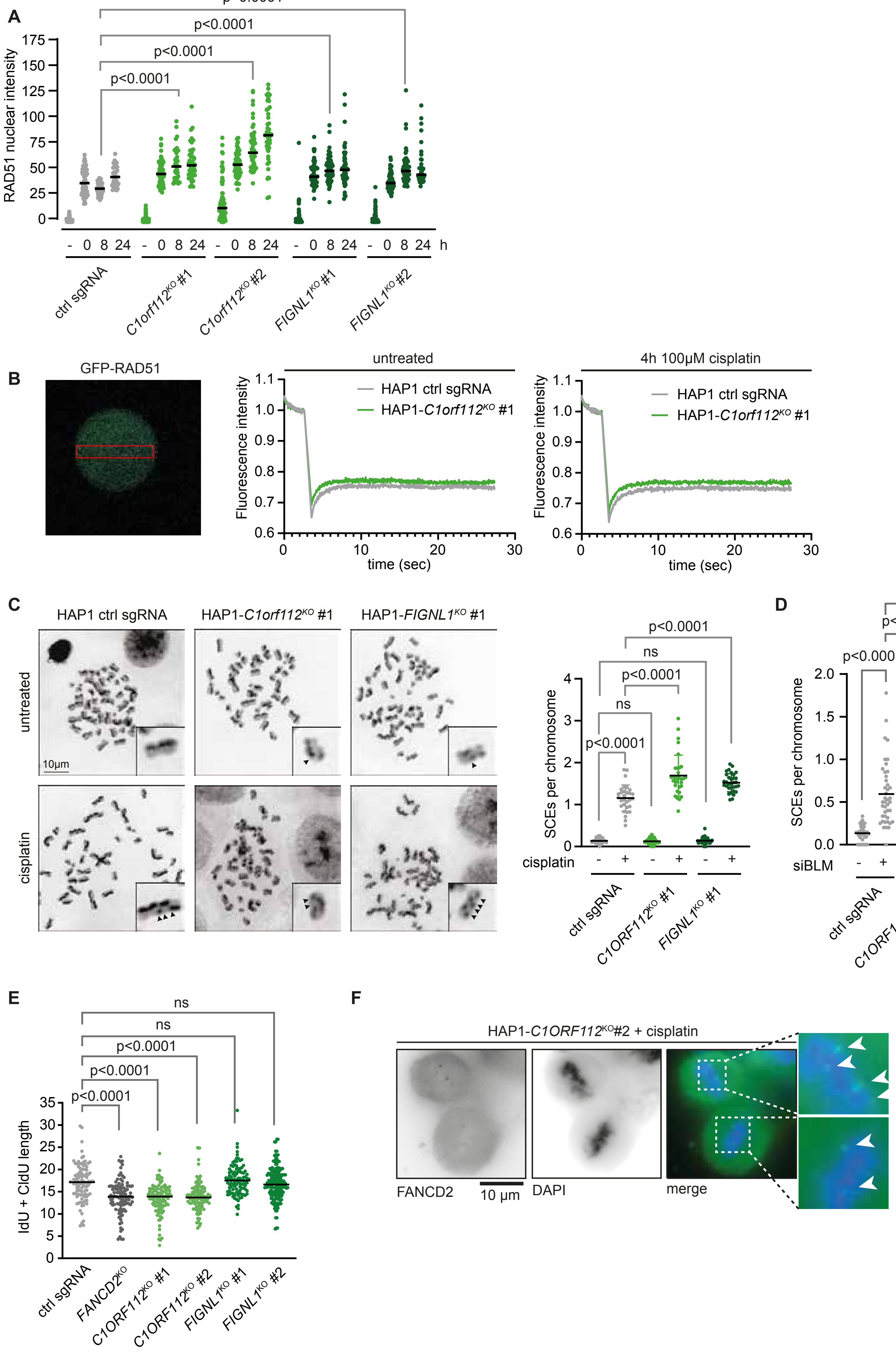

### Figure S5

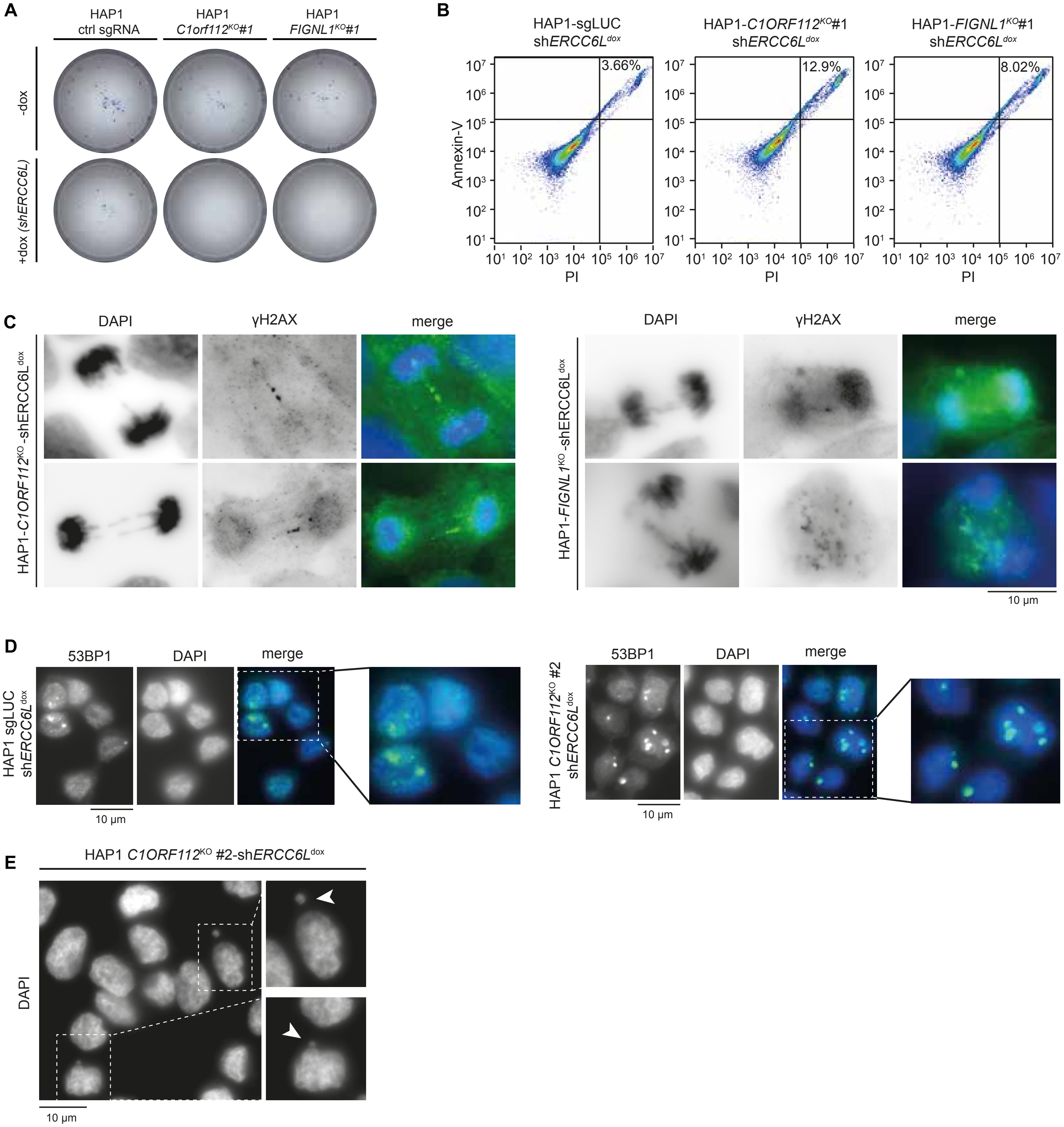
